# Temporal dynamics of RNA phage infection reveal bacterial recovery despite ongoing viral replication

**DOI:** 10.64898/2026.10.19.760031

**Authors:** Noam Harel, Yuval Resisi, Moran Meir, Nufar Buchshtab, Debjani Ghosh, Ido Golding, Uri Gophna, Adam Rosenthal, Adi Stern

## Abstract

Bacterial defenses against DNA phages are well documented, yet responses to RNA phages remain largely unexplored. Here, we characterize the population dynamics and host responses of *Escherichia coli* infected by the lytic RNA phage MS2 across multiple timescales, using bulk and single-cell transcriptomics, fluorescence microscopy, whole-genome sequencing, and genetic perturbations. Despite rapid viral takeover and extensive cell death, the bacterial population subsequently rebounds, revealing an unusual state in which host proliferation persists under intense phage pressure. Early in infection, heterogeneous subpopulations are evident, including one with low or undetectable viral transcript levels, low expression of the F-plasmid genes encoding the F-pilus (the MS2 receptor), and a distinct metabolic state. Strikingly, the rebounding population carries MS2 RNA in the great majority of cells, with evidence for active viral replication and continued cell division despite high viral loads. This rebound coincides with strong induction of biofilm-associated genes, including the *pga* operon, which synthesizes the biofilm-matrix exopolysaccharide PNAG. Deletion of *pgaA* causes earlier population decline and reduced population survival, indicating that *pga*-dependent matrix production delays viral lysis. Over longer timescales, phage-exposed populations show heterogeneous F-plasmid prevalence and plasmid-wide expression, consistent with dynamic cycling between plasmid-bearing and plasmid-free states: MS2 selects against F-plasmid-bearing cells, whereas conjugation may restore plasmid carriage. We propose that the biofilm-associated matrix delays viral lysis relative to continued cell division, long enough for rare F-plasmid-free daughter cells to arise and expand. Our results suggest that bacterial survival is determined by a dynamic race between viral lysis and host-cell division.

## Introduction

Bacteria have evolved a diverse arsenal of mechanisms that protect them from phage infection [1–3]. These defenses act at multiple stages of the viral life cycle, including adsorption, genome entry, genome replication, and infected-cell survival [4]. Many of the best-characterized systems target DNA phages, including restriction-modification [5], CRISPR-Cas [6], abortive infection [7], and recently discovered systems [8,9] that sense phage DNA [10], phage proteins [11], or infection-associated molecular signals [12]. In parallel, bacteria can also evade infection through physiological or structural changes, including receptor loss, receptor masking, altered surface composition, and biofilm-associated protection [4,13–15].

Compared with this expanding understanding of defense against DNA phages, bacterial responses against RNA phages remain much less explored. This gap is striking because RNA phages are no longer viewed as a narrow group of historical model systems [16,17]. Recent metatranscriptomic and meta-omic surveys have greatly expanded the known diversity of RNA phages [18–20], suggesting that they are widespread and ecologically important [21]. Yet the mechanisms that shape bacterial survival and population dynamics during RNA phage infection are still only beginning to be understood [18]. Only a handful of systems have been implicated in defense against RNA phages, including RNA-targeting CRISPR-Cas activities [22,23], toxin-antitoxin modules [24] and antiviral RNA ribonucleases [25,26].

The bacteriophage MS2, a positive-sense single-stranded RNA (+ssRNA) phage that infects *Escherichia coli*, provides a tractable model for studying dynamics of bacterial infection by an RNA virus. MS2 initiates infection by binding to the F-pilus [27], which is encoded by the *tra* operon [28]. The latter can be encoded on an autonomous plasmid or on the chromosome. When it is carried as a plasmid, the simplest route to escape infection is loss of the receptor, through plasmid loss or repression of pilus expression, and phage selection against F-plasmid carriage has been documented over longer timescales [29,30]. Whether receptor loss is the whole story, however, and how bacterial populations survive while most cells are already infected, remains unclear.

Addressing this requires resolving infection at the level of individual cells. Phage infection is inherently heterogeneous at the cellular level [31]: even within genetically identical bacterial populations, individual cells may differ in receptor availability [32], physiological state [33], infection timing [34], and the outcome of viral replication [35]. Such heterogeneity can be obscured by bulk measurements, which average infected, uninfected, and differentially responding cells. Recent advances in bacterial single-cell approaches, including single-cell transcriptomics [36–39] and single-molecule fluorescence in situ hybridization (smFISH) [40], now enable infection states and host responses to be characterized at cellular resolution, revealing rare subpopulations, intracellular viral burden, and cell-to-cell variation that are inaccessible to population-level profiling [37,39,41,42].

Here, we used MS2 infection of *E. coli* C-3000, a derivative of the K-12 strain that bears an F-plasmid, to examine how bacterial survival unfolds across cellular and population scales. Combining infection kinetics, bulk RNA-seq, single-cell assays, deletion strains, and whole-genome sequencing, we show that the bacterial population can recover even while most cells remain infected and viral replication continues. Infected cells continue to divide, while biofilm-related surface remodeling, particularly the *pgaABCD* pathway, delays phage lysis, thereby leading to increased survival. Over longer timescales, phage-exposed populations become heterogeneous in F-plasmid abundance and expression, consistent with selection for reduced receptor availability. We propose that the delayed lysis outcome afforded by surface remodeling creates time for daughter cells with lower viral burden and/or reduced receptor availability to arise and expand. Together, these findings reveal how changes in the timing of the viral replication cycle and in host population structure can reshape the outcome of RNA phage infection.

## Results

### Phage takeover is followed by population recovery despite ongoing viral replication

To obtain a view of host response to MS2 infection, we profiled viable host population (Colony Forming Units, CFU) alongside phage production (Plaque Forming Units, PFU) over time following phage infection at a high MOI of 10. At the high MOI employed, Poisson statistics predict widespread infection and cell death, with less than 0.005% of cells escaping infection [43]. Cell counts did decline sharply, but survival remained at roughly 2% of the uninfected control at 120 minutes (Fig. 1A, Methods), far above the Poisson prediction. This occurred even as phage titers exceeded 10¹¹ PFU/mL and outnumbered survivors by roughly four orders of magnitude. By 180 minutes the population had begun to rebound (Fig. 1A).

**Figure 1.**
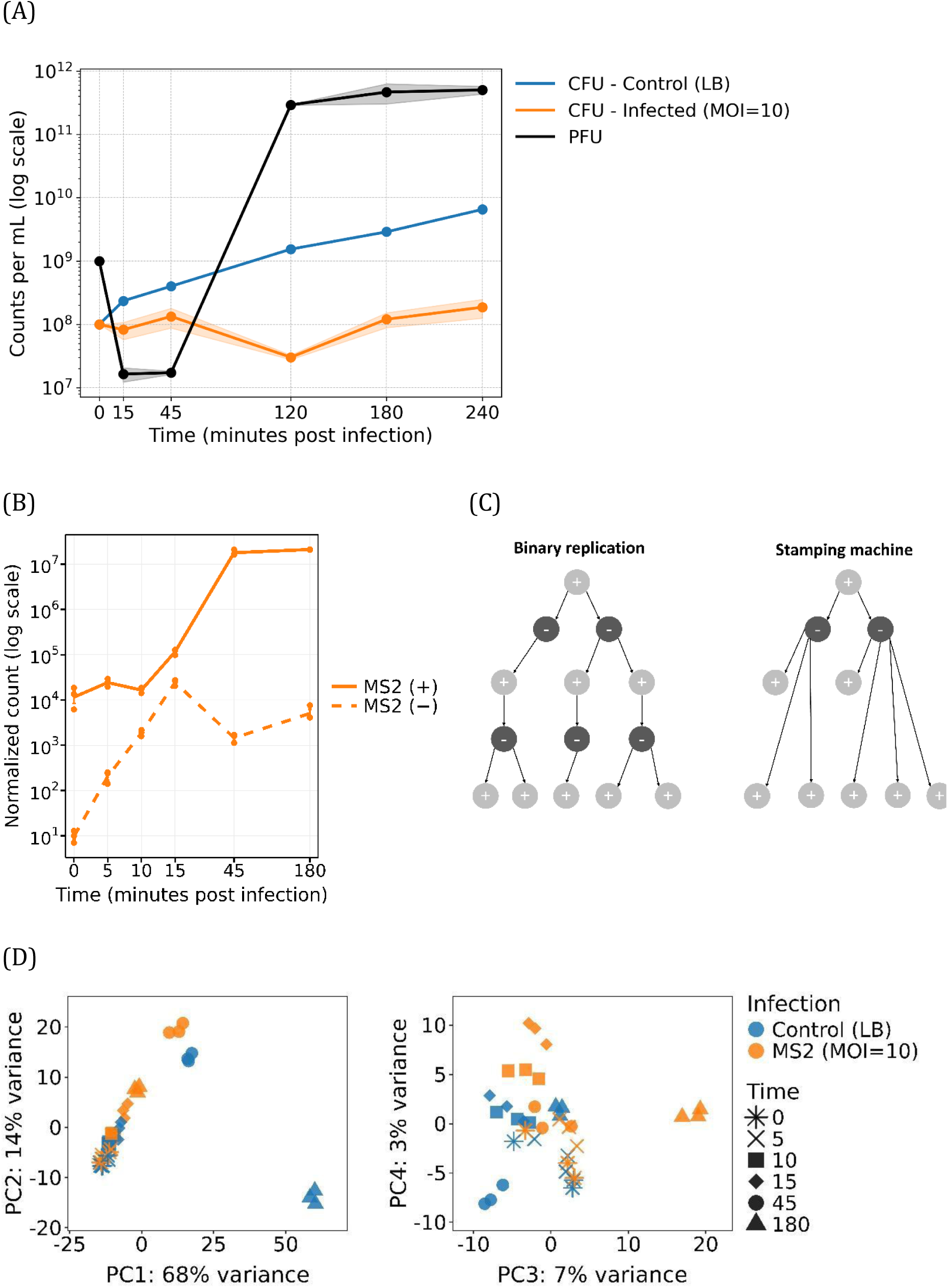
Population, viral RNA, and host transcriptome dynamics during MS2 infection. (A) *E. coli* C-3000 cell counts and MS2 phage titers following infection at an MOI of 10. Viable bacteria were quantified as colony forming units (CFU) in infected (orange) and uninfected (blue) cultures, and infectious phage were quantified as plaque forming units (PFU, black) in the cell-free culture supernatant. Values at time 0 represent the input at the time of infection: CFU were estimated from the optical density of the culture (*OD*_600_ of 0.5 ≈ 10^8^ cells/mL), and PFU from the titer of the phage stock. Curves represent the mean of two biological replicates for infected culture CFU and for PFU (shaded areas, ± SD); the uninfected control represents a single culture. (B) Strand-specific normalized MS2 RNA counts across the infection time course in MS2-infected cultures (MOI = 10), measured by bulk RNA-seq. Positive-strand (solid line) and negative-strand (dashed line) counts are plotted on a logarithmic y axis. Points represent biological replicates (n=3). (C) Binary versus stamping-machine modes of +ssRNA virus replication. Binary replication predicts parallel accumulation of both strands; stamping machine predicts early, transient negative-strand synthesis followed by positive-strand expansion, as observed in (B). Dark, negative strands; light, positive strands. (D) Principal component analysis (PCA) of bulk RNA-seq samples based on gene expression. Each point represents a biological replicate; colors indicate infection condition and symbols indicate time points. Left: PC1 and PC2, which account for 68% and 14% of the variance, respectively. Right: PC3 and PC4, which account for 7% and 3% of the variance, respectively. PC3 reveals infection-associated differences at 180 minutes that are less apparent in the first two principal components.

We set out to characterize host responses associated with population survival by using a combination of bulk RNA sequencing and single-cell profiling across different time points of the infection. To select time points spanning early infection through population recovery, we initially performed plaque assays and RNA quantitation (Fig. 1A, Fig. S1). This revealed that RNA replication began already by 10 minutes-post infection, and that by about 180 minutes the number of plaque forming units (PFUs) plateaued. Based on these kinetics, we performed bulk RNA sequencing across six time points (post-infection: 0, 5, 10, 15, 45, and 180 minutes) alongside uninfected control cells at parallel times, using three biological replicates per condition to yield a comprehensive set of 36 samples. To complement these bulk measurements, we additionally conducted single-cell RNA sequencing (scRNAseq) and smFISH, with uninfected controls for selected timepoints (Methods).

Unexpectedly, population recovery at 180 minutes occurred while intracellular viral RNA remained abundant. Positive-strand MS2 RNA continued to accumulate, and negative-strand RNA, indicative of viral replication, was still detectable at 180 minutes (Fig. 1B). Thus, the recovering population did not simply appear to consist of cells that had escaped viral exposure, raising the possibility that host growth and viral replication were occurring concurrently.

### Strand-specific RNA dynamics reveal distinct phases of MS2 replication

Strand-specific viral RNA dynamics revealed an early increase in negative-strand RNA genomes, which peaked at approximately 15 minutes and then rapidly declined, while positive-strand RNA genomes underwent massive expansion between 15 and 45 minutes (Fig. 1B). At early time points, negative-strand reads were strongly enriched toward the 3′ end of the MS2 genome (Fig. S2), consistent with accumulation of newly initiated negative strands, although differential RNA degradation could also contribute to this pattern. Together, these dynamics are consistent with a stamping-machine mode of replication, in which negative-strand templates are generated early and subsequently support extensive positive-strand production [44] (Fig. 1C).

### Minimal alterations to host transcription during early stages of infection

We analyzed the bulk RNA-seq host transcriptome using two comparisons: infected versus uninfected samples at matched time points, and successive time points within each condition. We observed minimal differential expression between infected and uninfected samples during the period of 0 to 15 minutes, both when comparing infected to uninfected samples and across these early time points (Fig. 1D).

The number of genes differentially expressed between infected and control samples increased at 45 minutes relative to any earlier time point (Fig. S3, Table S1). This number was nonetheless smaller than the number of genes differentially expressed between infected samples at 45 and 15 minutes, indicating that progression of the culture, rather than infection, was the dominant source of transcriptional change at this stage. Consistent with this, infected and uninfected 45 minute samples clustered together, apart from the earlier time points (Fig. 1D). At 180 minutes post-infection, a much more pronounced shift in host transcriptomic profiles was evident in infected compared to uninfected samples. Along the major principal components, the 180 minute infected samples clustered closer to the early (≤15 minute) samples than to the corresponding 180 minute controls (Fig. 1D). We reasoned that this shift could partly reflect the extensive progressive cell death, which reduced population density and effectively returned the culture to an early-growth-like state. We revisit the 180 minute time point below.

Given the rapid replication of the virus already by ten minutes, the minimal response of the host to MS2 was surprising. We searched for genes that were upregulated across multiple timepoints within the first 15 minutes of infection. This revealed a total of ten such genes (Fig. S4). Of these, two stood out as related: *nhaR*, a transcriptional regulator [45], and the gene regulated by it *nhaA*, an Na+/H+ antiporter, which we will discuss later.

### Single-cell analyses reveal an early low-MS2 subpopulation with a distinct host state

To capture host-phage dynamics at cellular resolution, we performed scRNAseq profiling using the ProBacSeq method [36] (Methods). Briefly, this approach relies on cell permeabilization followed by the introduction of specialized probes designed to hybridize to host mRNAs (and in our setup, MS2 viral RNA) while encoding poly-A tails and unique molecular identifiers (UMIs) for downstream sequencing on the 10x Genomics platform. Our probe set comprised targeted *E. coli* gene probes supplemented with additional probes specific to both the forward and reverse strands of the MS2 bacteriophage.

scRNAseq pseudo-bulk expression correlated strongly with matched bulk RNA-seq (Pearson R^2^=0.71, p<10^-10^), with anomalously captured host genes excluded from downstream analyses (Fig. S5). At 15 minutes post-infection, viral signal was heterogeneous across the population and close to the detection limits needed to confidently assign individual cells to infected and uninfected states. We therefore used this time point primarily to assess early heterogeneity and focused the host-transcriptional analysis on 30 and 45 minutes post-infection, when infection state classification was more robust (Fig. S6).

At both 30 and 45 minutes, we detected a distinct subpopulation of cells with low or undetectable MS2 signal, comprising ∼4–10% of the cellular population (Fig. 2A, S7). We denote this subpopulation as exposed-uninfected. Two additional subpopulations at 45 minutes showed high *fim* operon expression and high *srl* operon expression, respectively, and were observed both in infected and control populations (Fig. S8). We therefore excluded these operons from downstream analyses. Notably, type I fimbriae undergo phase variation through inversion of the *fimS* promoter [46], and we attributed the observed heterogeneity in the *fim* operon to constitutive phase switching. This interpretation was further supported by *fimS* inversion in the genomic data described below (Table S5).

**Figure 2.**
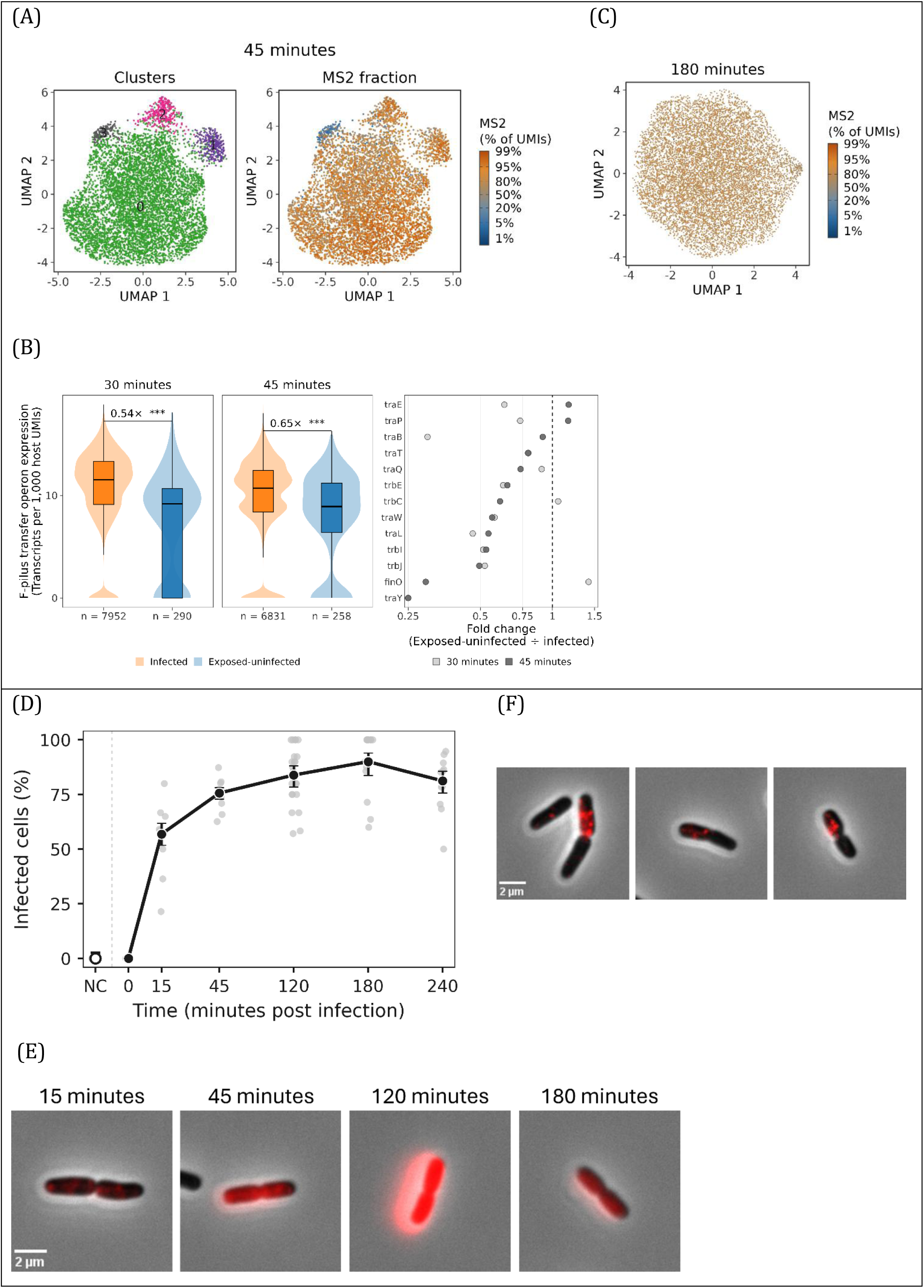
Single-cell dynamics of MS2 infection and host transcriptional states. (A) UMAP visualizations of scRNAseq expression data at 45 minutes post-infection. Left: Cells are colored by cluster identity (0-3) based on clustering of host transcripts. Right: Cells are overlayed by the continuous percentage of MS2 viral transcripts. Cluster 1 is characterized by *fimA* upregulation, and Cluster 3 represents an exposed-uninfected population, characterized by higher levels of *glp/mal* regulons and low/absent levels of MS2. (B) Left: Per-cell expression of the F-pilus transfer operon, as transcripts per 1,000 host UMIs (log scale), in infected (orange) vs exposed-uninfected (blue) cells at 30 and 45 minutes. Violins show the full distribution; boxes mark median and IQR. Fold change and significance from a negative-binomial GLM with host-UMI offset; ***P < 0.001. Right: Fold change (exposed-uninfected ÷ exposed-infected) for individual well-detected transfer-region genes at 30 minutes (light grey) and 45 minutes (dark grey); dashed line marks no change. (C) 180 minutes scRNAseq shows vast majority of cells infected. (D) Percentage of MS2-positive cells measured by smFISH across infection time points. Grey points represent individual microscopy images and black circles indicate pooled percentages, with error bars showing Wilson 95% confidence intervals. Infection was defined using a positive-strand MS2 signal threshold set above the maximum observed in the negative control (NC, open circle). (E) Representative smFISH images of MS2-positive cells undergoing division along time points. (F) In three dividing pairs at 15 minutes post-infection, MS2 signal is detected more prominently in one daughter cell, illustrating asymmetric partitioning of viral RNA during cell division.

Differential expression analysis first revealed that the exposed-uninfected cells had reduced expression of F-plasmid *tra* operon genes (negative-binomial GLM with host-UMI offset, P < 10⁻¹⁰, Fig. 2B). However, plasmid-derived transcripts remained detectable, and the sparse single-cell signal prevented us from distinguishing between partial plasmid loss and transcriptional downregulation of plasmid genes. The exposed-uninfected subpopulation also showed higher expression of the *mal* (maltose) and *glp* (glycerol) regulons, which emerged by 45 minutes. We found that these cells, unlike infected and control cells, did not reduce *mal*/*glp* expression (Fig. S9). This signature may reflect scavenging of nutrients released by lysing neighbors [47] or carbon limitation, as both regulons are catabolite-repressed and cAMP-CRP-dependent [48]; in either case, we postulate that this metabolic response is unlikely to be a major driver of population recovery.

Analysis of the global host transcriptome at this 45 minute milestone provided a broader view of the population-wide response. Functional enrichment of host genes upregulated in the bulk RNA-seq data at 45 minutes post-infection revealed a prominent induction of *de novo* UMP biosynthesis and pyrimidine metabolism pathways (Table S2). This population-wide signature was not associated with any distinct subset of cells in scRNAseq analysis and is consistent with increased ribonucleotide demand imposed by extensive viral RNA replication [49].

### Population recovery occurs despite widespread infection and continued cell division

We next examined scRNAseq at 180 minutes post-infection. In contrast to 45 minutes, we did not detect distinct host-transcriptional subpopulations. All detected cells at 180 minutes contained substantial MS2 RNA (median ∼80% of total transcripts) (Fig. 2C). Thus, the late population appeared markedly more homogeneous with respect to the host transcriptional states that could be resolved, although the dominance of viral transcripts limited sensitivity for detecting subtler differences.

We independently assessed infection prevalence by smFISH at corresponding infection time points, with probes targeting positive-sense MS2 [50] (Methods). Approximately 57% of cells were MS2-positive at 15 minutes, increasing to 76–81% at 45 minutes across technical replicates and exceeding 80% from 120 minutes onwards (Figs. 2D, S10, Table S3, Methods). These values fall somewhat below the ∼90–96% obtained by scRNAseq, which excludes low-RNA cells that smFISH retains and scores as negative (Methods, Fig. S7). Thus, both scRNAseq and smFISH reiterated the bulk RNAseq results (Fig. 1B), whereby population recovery at 180 minutes occurred despite widespread infection.

Across all time-points profiled by smFISH we observed MS2-positive cells undergoing cell division, showing that cells carrying intracellular viral RNA can continue to divide [51] (Fig. 2E). Anecdotally, in three dividing cell pairs at 15 minutes, MS2 signal was detected more strongly in one daughter cell (Fig. 2F). This supports unequal partitioning of viral RNA during division and the potential generation of a daughter cell with little or no detectable viral burden [52,53]. Although independent or later infection of only one daughter cannot be excluded, the cells were captured during septation, leaving only a short interval in which such differential infection could have occurred.

### Population recovery is accompanied by cell-surface remodeling

We next analyzed differential gene expression at the later time point of 180 minutes post-infection. Notably, bacterial counts (CFU) rebounded at this time point while viral titers (PFU) remained high and stable (Fig. 1A). As described above, four observations indicate that viral RNA remained abundant and replication was ongoing at this time point: positive-strand MS2 RNA was still abundant and negative-strand replication intermediates were still present at high levels (Fig. 1B); MS2 transcripts made up a median of ∼80% of the transcriptome per cell by scRNAseq (Fig. 2C); and ∼90% of cells were MS2-positive by smFISH (Fig. 2D). Together with the rebound in bacterial counts, this indicates a population that was expanding while under substantial phage pressure, both intracellular and extracellular. Unlike the earlier time points, we observed a striking transcriptomic shift at this stage, with hundreds of genes, encompassing approximately one-quarter of the genome’s coding regions, showing significant differential expression compared to the 180 minutes control time point (Fig. 1D, Fig. S3). Because the matched 180 minutes control also differed substantially from earlier controls, this shift could include both infection-specific and growth-phase effects (Fig. 1D).

To enrich for infection-specific changes, we compared 180 and 15 minute infected samples and excluded genes that showed comparable changes between the corresponding control time points. The remaining upregulated genes were enriched for cell-surface remodeling and biofilm-related processes, including slime-layer polysaccharide biosynthesis, colanic acid metabolism, and cell adhesion [15,54,55] (Fig. S3, Table S4). Particularly prominent was induction of the *pgaABCD* exopolysaccharide system [56], a known target of *nhaR* [45], which was itself strongly upregulated across all the early infection time points (Fig. S4). This temporal pattern aligns with the observed gene expression dynamics: *nhaR* exhibited a sharp increase in expression within the first few minutes of the experiment, while *pgaABCD* began increasing at 45 minutes, eventually reaching high expression levels alongside other surface-remodeling genes such as the *wca* colanic acid metabolism operon genes [57,58] in the 180 minute post-infection population (Fig. 3A).

**Figure 3.**
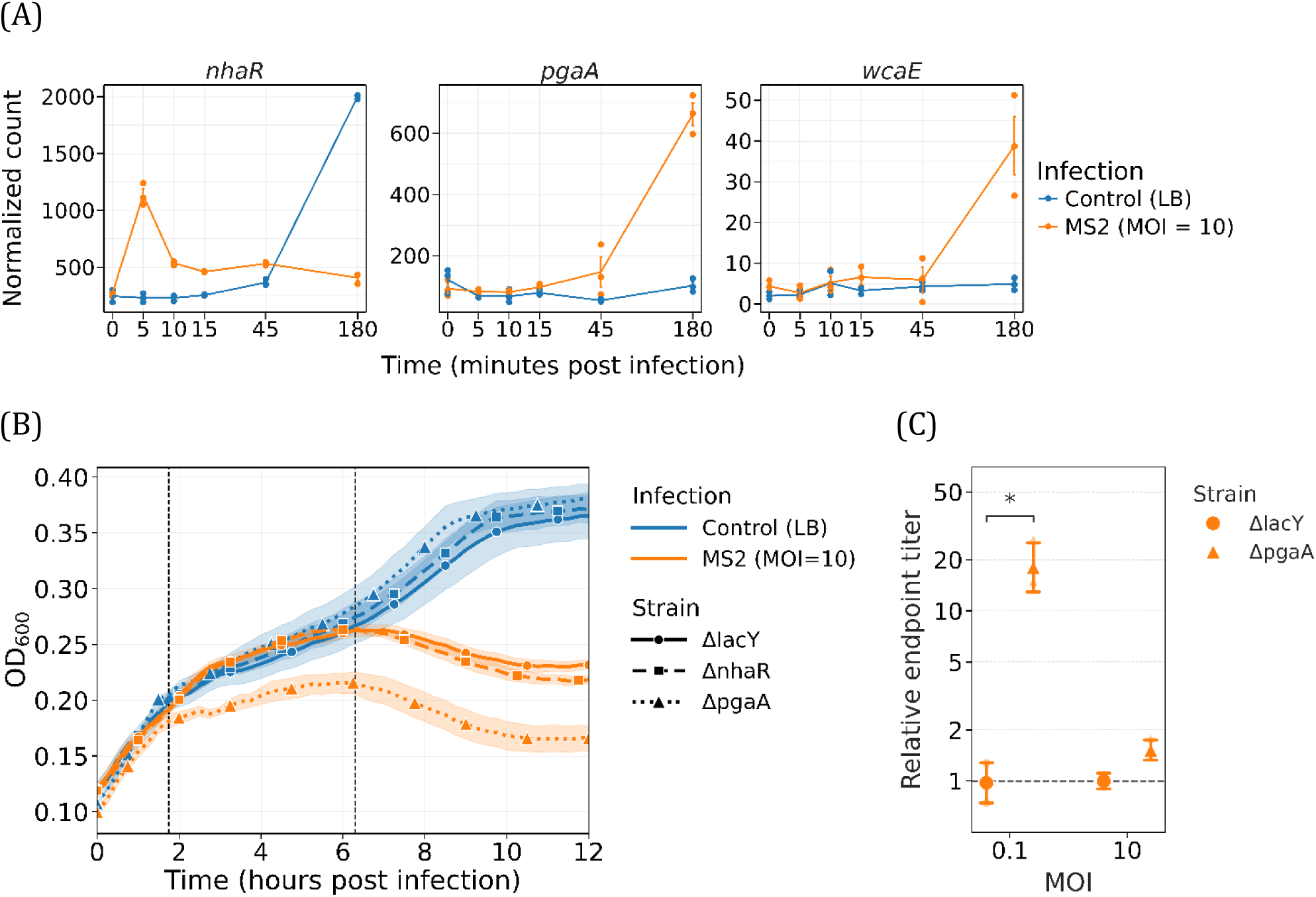
Expression dynamics and knockout phenotypes implicate *pgaA* in protection against MS2 by delaying lysis. (A) Change in expression over time of genes *nhaR*, *pgaA,* and *wcaE*, plotted as normalized counts. (B) Liquid culture growth of *E. coli* strains with deletions of *lacY* versus a deletion of *nhaR* and *pgaA*. Cells were infected by MS2 at an MOI of 10 (orange) or left uninfected (blue). Measurements were performed at *OD*_600_, in three biological replicates; lines show the mean and shaded areas the SD. Dashed vertical lines indicate the onset of divergence from the corresponding uninfected control, for Δ*pgaA* (left) and Δ*lacY*, Δ*nhaR* (right). (C) Relative endpoint titer, defined here as the PFU titer recovered from the *ΔpgaA* strain after 3 hours of infection divided by the PFU titer recovered from the Δ*lacY* control strain under the same conditions, at MOI 0.1 and 10. Large markers show the mean ± SD of biological replicates (n = 3, except Δ*lacY* at MOI 10, n = 2), and small markers the individual replicates. * two-sided t-test, p < 0.05; the difference at MOI 10 was not significant (p = 0.052).

### Surface-associated genes contribute to protection against MS2

To test whether surface-associated pathways contribute to protection against MS2 phage, we used knockout strains with deletions of *nhaR* and *pgaA*, with a *lacY* deletion strain as a control [59] (Methods). We infected each strain with MS2 at an MOI of 10 in liquid culture and compared cell density to mock infections. Deletion of *pgaA* produced both an earlier decline in cell density and a lower surviving population than the Δ*lacY* control following MS2 infection (Fig. 3B). Notably, the divergence from the uninfected control, which we use to estimate the onset of major lysis (dashed lines, Fig. 3B), occurred earlier in *ΔpgaA* than in the control strains. This provides a timing-based indication that *pgaA* delays the onset of phage-induced population decline in addition to altering the extent of killing. The absence of a detectable phenotype in the *nhaR* knockout may reflect regulatory redundancy or incomplete dependence of the protective response on *nhaR,* which operates alongside post-transcriptional and allosteric control of the Pga machinery [60,61].

To further test whether *pgaA* provides direct protection against MS2, we measured endpoint phage titers in the *pgaA* knockout relative to the control. We observed a significantly higher relative endpoint titer in the *pgaA* knockout strain at a low multiplicity of infection (MOI = 0.1), resulting in a more than tenfold increase in viral titer relative to the control (p<0.05, t-test) (Fig. 3C). A similar increase in phage production was observed at a higher multiplicity of infection (MOI = 10), although this difference was of borderline significance (p = 0.052, t-test). The higher endpoint phage yield in Δ*pgaA* (Fig. 3C) is consistent with a role for *pgaA* in delaying infection, as an earlier and more extensive population decline in the absence of *pgaA* permits more infection cycles within the assay window.

### Population recovery at 180 minutes is not explained by population-wide F-plasmid loss or pilus downregulation

MS2 initiates infection by binding specifically to the F-pilus, encoded by the *tra* operon on the F-plasmid [27,28]. We therefore examined the transcriptional dynamics of plasmid-borne genes, treating the *tra* operon separately from the remaining F-plasmid genes (Methods). At 45 minutes post-infection, *tra* operon expression was modestly lower in infected than in time-matched control cultures (0.77-fold, P = 0.008). The remaining plasmid genes showed no such reduction and were, if anything, higher in infected cultures (1.31-fold, P = 0.007), indicating regulation of the *tra* operon specifically rather than loss of the plasmid. The reduction was nonetheless small and unevenly distributed across the operon, and did not include the gene encoding the pilin subunit itself, the most abundant transcript of the operon, which was unchanged (0.99-fold, P = 0.87; Fig. S11).

The *tra* operon was not progressively silenced as the population recovered; it was maintained. Between 45 and 180 minutes in infected cultures, expression of the *tra* operon and of the F-plasmid both changed only slightly (0.89- and 0.7-fold, respectively), and absolute module scores remained high throughout (Fig. 4A). The pilin gene itself declined modestly over this interval (0.59-fold, Fig. S11), although the operon as a whole was maintained. The larger apparent difference between infected and control samples at 180 minutes was driven by the controls, in which F-plasmid expression rose approximately twofold as the culture slowed (Fig. 4A). This was expected: F-plasmid copy number is known to rise as growth rate falls [62], and the controls, unlike the lysis-depleted infected cultures, had begun to approach stationary phase by this time (Fig. 1A). To summarize, bacterial counts rose at 180 minutes in the presence of more than 10¹¹ PFU/mL, with most cells still carrying MS2 RNA (Figs. 1A, 1B, 2C, 2D). Recovery therefore occurred with the receptor operon mostly intact, and was not achieved by population-wide receptor downregulation or plasmid loss.

**Figure 4.**
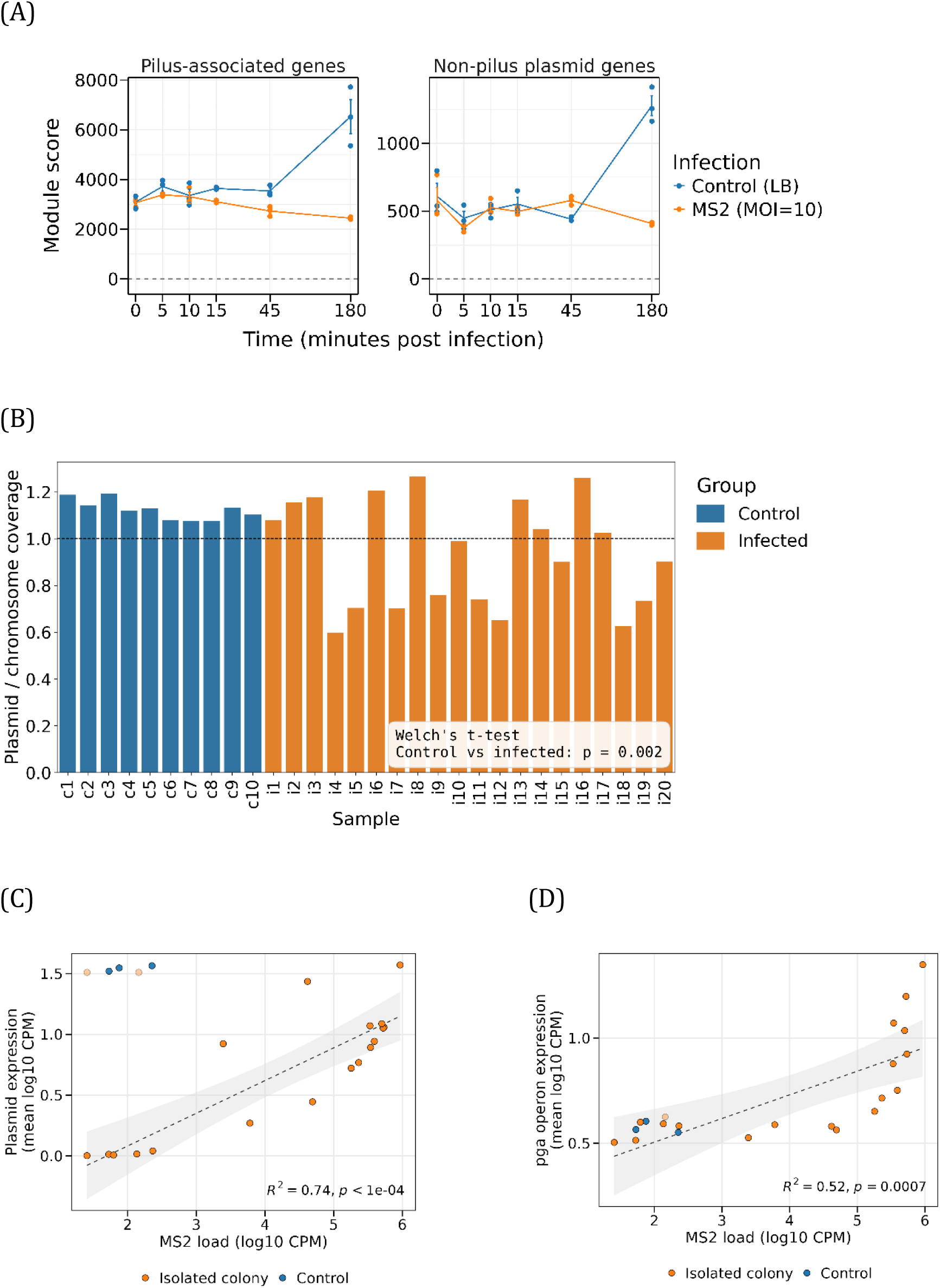
F-plasmid expression and abundance across acute and long-term MS2 exposure. Panel (A) reports the bulk RNA-seq time course over the first 180 minutes of infection (acute infection); panels (B–D) report colony isolates recovered after overnight phage exposure and regrowth. (A) Mean module scores (average normalized transcript counts) of the pilus-associated genes (n = 33) and non-pilus F-plasmid genes (n = 53) across the infection time course and corresponding uninfected controls. Points represent biological replicates. (B) F-plasmid-to-chromosome coverage ratios calculated from whole-genome sequencing of phage-exposed and unexposed colony isolates. (C) Mean module scores of F-plasmid genes (n = 79 expressed genes) plotted against MS2 load, defined as log-normalized positive-strand MS2 RNA counts (log10 CPM of host-mapped reads); module scores are the mean across member genes of log10(CPM + 1). Lines show linear regressions across the 18 plaque-negative isolates with 95% confidence bands. Control samples (blue) and two colony-derived revertants (light orange) were excluded from the regressions. (D) As in (C), for the *pga* operon. Regressions, confidence bands and excluded samples in (D) as in (C). A corresponding analysis with F-plasmid genes partitioned into pilus-associated and non-pilus sets is shown in Fig. S12.

### Most surviving isolates carry no fixed genetic mutations

We next asked whether selected mutations could account for the rapid rebound in CFU at 180 minutes or for the exposed-uninfected subpopulation at 45 minutes. Because cells from these early time points cannot be cloned without further growth, we sequenced the genomes of 20 colonies isolated from phage-exposed cultures and 10 unexposed control colonies (Methods). These colonies grew overnight on phage-containing plates and were re-streaked and regrown before DNA extraction, giving them considerably more opportunity to acquire resistance mutations, or to lose the F-plasmid, than cells in the acute time course. Mutations detected among them therefore represent an upper bound on those present during recovery: if mutations are rare even here, they are unlikely to explain events within the first 180 minutes.

We conducted long-read whole-genome sequencing on all 30 isolates using the Oxford Nanopore Technologies (ONT) platform. Genomic analysis revealed no fixed synonymous or nonsynonymous mutations across the 10 unexposed control isolates. Notably, in nearly all isolates, a fraction of reads (∼20–30%) supported inversion of the *fimS* promoter element, consistent with the phase-variable *fimA*-high subpopulation observed by scRNAseq (Table S5). Among the 20 phage-exposed isolates we identified only four fixed substitutions, each unique to a single isolate: one synonymous, one non-synonymous in *yejK*, encoding a nucleoid-associated protein [63], and two non-synonymous in structural genes of the colanic acid (*wca*) biosynthesis operon [57]. This pathway has been implicated in phage resistance through increased exopolysaccharide production and reduced phage adsorption [4,64,65], and we found it upregulated at 180 minutes post-infection (Fig. 3A). Because no genomic changes were found in 16/20 phage-exposed isolates (Table S5), we concluded that fixed genetic mutations do not account for the survival phenotype of most of them, pointing instead toward other mechanisms.

To investigate alternative genomic factors, we next quantified the F-plasmid-to-chromosome copy number ratio across all samples. This analysis revealed a distinct structural pattern: while a subset of the phage-exposed isolates maintained plasmid-to-chromosome ratios comparable to the unexposed controls, several other phage-exposed isolates exhibited a much lower ratio (Fig. 4B). Given the low copy number of the F-plasmid (∼1–2 copies per cell) [66], this pattern is consistent with partial plasmid loss within the population grown from each isolate. Reduced F-plasmid abundance would be expected to reduce F-pilus availability and thereby MS2 susceptibility [30]. Given the scarcity of fixed genomic mutations and the heterogeneous plasmid abundance observed among the isolates, we set out to investigate the transcriptomic profiles of these isolates.

### Phage-surviving colonies retain dynamic heterogeneity in MS2 load and F-plasmid expression

From the 20 isolate stocks used for genomic analysis, we selected 12 for bulk RNA-seq, including four isolates profiled with biological replicates to assess within-isolate transcriptomic variation, and three uninfected control replicates (Table S6). Bulk RNA-seq revealed substantial heterogeneity among the phage-exposed isolates. In contrast to the uninfected controls, which clustered tightly, phage-exposed isolate replicates showed divergent transcriptional profiles, indicating that the post-exposure phenotype was associated with heterogeneous population states rather than a uniform expression program (Fig. S13). To identify the biological factors driving this heterogeneity, we examined the transcripts contributing most strongly to variation across samples. The dominant variable features included MS2 positive- and negative-strand transcripts, together with genes distributed across the F-plasmid. Several samples retained high levels of viral RNA despite repeated isolation and subculturing before RNA extraction. The presence of MS2 negative-strand reads indicated ongoing viral replication [67], suggesting that these phage-surviving colony-derived populations remained heterogeneous and contained actively infected cells. We also considered whether changes in the phage population itself might have contributed to the observed dynamics, since phage populations can accumulate defective or otherwise aberrant genomes during replication [68,69]. We therefore examined the MS2-derived RNA-seq reads for viral sequence variants. Although a small number of mutations were detected (data not shown), we found no evidence of a distinct or defective viral genotype expanding within the population.

We next asked whether variation in MS2 abundance reflected variation in F-plasmid state. Plasmid-derived expression varied by more than three orders of magnitude across isolates, from near-complete loss to control-level expression, and MS2 abundance tracked this variation closely (Fig. 4C; *R*² = 0.74, *p* = 5.3 × 10^-6^). Two isolates departed from this relationship, carrying F-plasmid transcripts at control levels yet with little or no detectable MS2. To test whether these populations remained susceptible, we challenged all isolates with phage in a plaque assay. These two isolates were the only ones to support plaque formation; all others were plaque-negative. Both had therefore cleared the virus and reverted to a susceptible, uninfected state rather than acquiring resistance. Because they occupy a biological state distinct from that of the phage-surviving isolates, they were excluded, together with the unexposed controls, from the regressions reported here.

To ask whether this association was specific to the MS2 receptor, we partitioned the plasmid into the pilus-associated genes encoding the conjugative pilus and all remaining plasmid genes (Fig. S12). Thus, the major axis of variation was not restricted to receptor-gene expression; rather, viral load tracked plasmid-wide expression, consistent with variation in the fraction of F-plasmid-bearing cells within each colony-derived population.

We further tested whether any of the biofilm-related genes were associated with MS2. This analysis revealed that the *pgaABCD* operon was correlated with MS2 across the isolates. The four *pga* genes behaved as a single transcriptional unit across isolates (mean pairwise *r* = 0.88), and their combined expression scaled with MS2 abundance (*R*² = 0.52, *p* = 7.3 × 10⁻⁴; Fig. 4D).

Together, these results distinguish two timescales of population survival. Acute recovery occurs despite widespread intracellular MS2 and coincides with induction of the *pga* biofilm machinery, pointing to a rapid, physiological route to protection.

Longer-term phage exposure is associated with heterogeneous F-plasmid states and receptor availability rather than recurrent fixed chromosomal mutations.

## Discussion

Our results reveal an unusual response to MS2 infection that is not well described by classical resistance, in which cells simply prevent or terminate infection [1,3,4]. Instead, bacterial population recovery occurs despite continued high viral burden. At 180 minutes, when CFU had begun to rebound, positive-strand viral RNA remained abundant, negative-strand replication intermediates were still detectable, and both scRNAseq and smFISH showed MS2 RNA in the great majority of cells. Moreover, smFISH captured MS2-positive cells undergoing division. Together with the phenotype inferred via the *pgaA* pathway, these observations suggest a model in which the outcome of infection reflects a changing balance between phage-induced cell death and continued host-cell proliferation (Fig. 5). Longer-lived associations in which bacteria continue to grow while propagating phage, have been described as carrier states or chronic infections [70], including for a single-stranded RNA phage [71].

**Figure 5.**
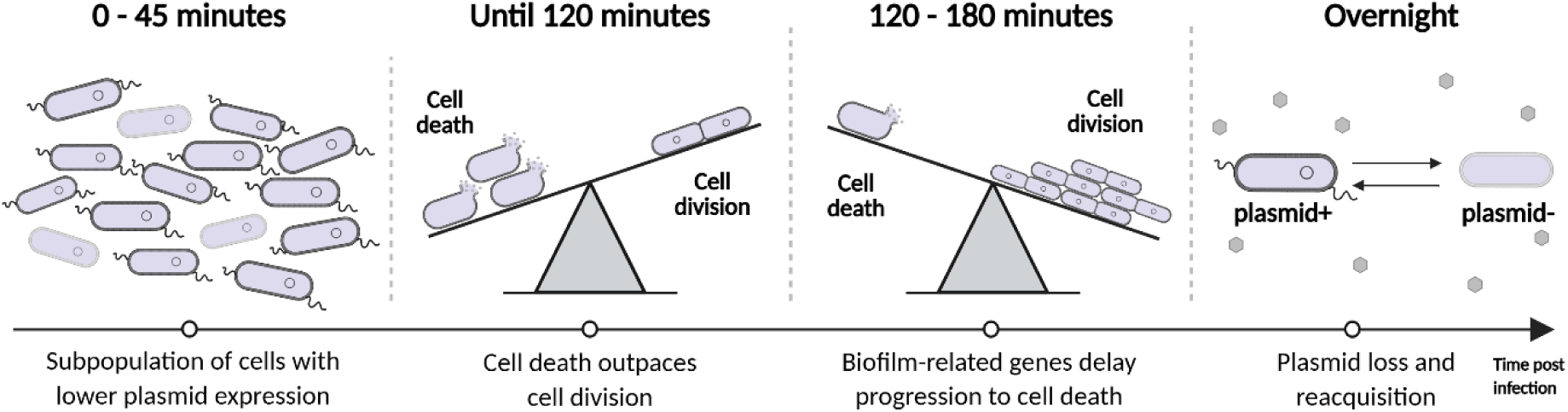
Proposed model of population survival and recovery during MS2 infection. Early infection produces heterogeneous cellular outcomes, followed by extensive lysis. Induction of surface-remodeling pathways, including *pgaABCD*, is proposed to delay progression to lysis and shift the population-level balance toward continued cell division despite widespread viral infection. Over longer timescales, MS2 imposes negative frequency-dependent selection on F-plasmid carriage. When plasmid-bearing cells are common, viral load is high and selection strongly favors plasmid loss; as these cells become rare, viral pressure relaxes, allowing reacquisition of the plasmid through conjugation. Created in BioRender. Resisi, Y. (2026) https://BioRender.com/2e8793w

### Early Subpopulation Heterogeneity

scRNAseq at 45 minutes post-infection resolved a distinct subpopulation, comprising ∼4–10% of host cells, characterized by low or undetectable MS2 transcripts and reduced expression of the F-plasmid *tra* operon, which encodes the MS2 receptor. We interpret this as a transient state of low receptor availability, since plasmid transcripts remained detectable and stochastic variation in receptor and conjugative-pilus expression is a known route by which sensitive cells transiently evade adsorption [32,72]. Some of these cells may therefore simply be on their way to infection, delayed rather than spared. Their retained *mal* and *glp* expression is more plausibly a consequence of remaining uninfected than a cause of it (Fig. S9).

### Surface Remodeling Shifts the Balance Between Lysis and Cell Division

A second phase emerges as the population transitions from extensive cell loss toward recovery. This transition coincides with strong induction of surface-associated, biofilm-related pathways, including *pgaABCD*. Although K-12 strains are relatively weak biofilm producers under standard laboratory conditions [73], these pathways can alter the bacterial surface independently of mature biofilm formation. Importantly, deletion of *pgaA* caused both an earlier decline in cell density and a lower surviving population following MS2 infection. The timing of this phenotype argues that *pgaA* does not act simply by determining whether infection succeeds, but by altering its kinetics - delaying the interval between infection and lysis. Consistent with a broader role for biofilm-related surface remodeling, two independently derived phage-surviving isolates carried fixed nonsynonymous mutations in the colanic-acid biosynthesis genes, which we also observed were upregulated at 180 minutes post-infection (Fig. 3A). Colanic-acid production has previously been linked to phage resistance in *E. coli*, including through receptor masking and reduced adsorption [64,65]. Although these mutations were too rare to explain population recovery broadly, their recurrence within the same pathway provides independent genetic support for the involvement of biofilm-related surface pathways during MS2 exposure. The phage-surviving isolates point the same way, with *pgaABCD* expression scaling positively with MS2 abundance, consistent with the surface program remaining engaged as long as infection persists without clearing it.

We therefore propose that the surface response alters the balance between lysis and continued host proliferation (Fig. 5). Early after infection, the rate of lysis exceeds that of cell division and the bacterial population contracts. By approximately 180 minutes, however, host proliferation becomes sufficient to offset or exceed continuing cell loss, allowing population recovery even though viral replication persists. *pgaA* provides the clearest functional support for this model: its deletion shifts the dynamics toward earlier lysis, while other induced surface-remodeling pathways may contribute similarly [74].

The *pga*-dependent matrix need not act at a single step, and it may delay both entry and lysis. At an MOI of 10, the majority of cells are infected as we show herein; this isolates the post-entry delay but would mask any contribution at entry, since a barrier to incoming virions can be overcome by excess phage. Such a contribution may nonetheless exist: the phage-yield increase in Δ*pgaA* was pronounced at MOI 0.1 yet only marginal at MOI 10 (Fig. 3C). Future work will be required to elucidate the relative contribution of these two steps.

A delay in lysis could support population recovery through several routes. Infected cells may divide before lysis, with viral material diluted or asymmetrically partitioned between daughters, potentially producing cells with substantially lower viral burden [53]. Daughter cells may also enter physiological or receptor states that reduce subsequent infection or reinfection. In three dividing pairs imaged at 15 minutes, MS2 signal was more intense in one daughter than in the other (Fig. 2F). We offer this as an illustrative observation rather than a quantitative result: three pairs are anecdotal, and we cannot formally rule out that one daughter was infected independently or slightly later, rather than inheriting viral RNA unequally. That said, capture during septation leaves only a brief window for such differential infection. We also expect such asymmetric partitioning to be rare and short-lived and correspondingly hard to capture. Establishing whether asymmetric partitioning contributes meaningfully to population recovery will require dedicated, quantitative time-resolved imaging.

Delaying infection may represent an underappreciated form of protection in natural bacterial populations [75]. Rather than directly preventing infection, extending the lifetime of an infected cell could allow continued host division and thereby reduce the population-level cost of phage attack. Whether the host delays genome entry or extends the interval from entry to lysis, the consequence for the virus is the same: the time from encountering a host to releasing progeny is extended. Lysis timing is not a passive endpoint but a finely tuned phage trait. Theory predicts an optimal latent period that maximises phage fitness at a given host density [76,77], and experimental evolution of lysis time converges toward that optimum [78], although it does not always reach it. A host that extends any step of the cycle therefore displaces the phage from a timing optimum it is under selection to maintain, and even modest delays are expected to carry a fitness cost. More generally, delaying infection could provide time for less susceptible subpopulations to arise and expand, including plasmid-free cells with reduced receptor availability [30].

### The Tripartite Dynamic: Conflict Between Selfish Genetic Elements

Over longer timescales, the dominant selective pressure appears to shift toward the F-plasmid (Fig. 5). Phage-exposed isolates rarely carried fixed mutations, yet varied markedly in F-plasmid abundance and plasmid-wide expression, which closely tracked viral burden. Unlike conventional host-virus interactions, this system can therefore be viewed as a three player conflict among the bacterial chromosome, the F-plasmid, and MS2 [79]. The F-plasmid promotes its own transmission through the *tra* operon, while MS2 exploits the resulting F-pilus as its obligate receptor.

Our data is consistent with MS2 selecting for plasmid-free cells, thereby reducing F-pilus availability and viral entry. Because plasmid loss was incomplete, residual plasmid-bearing cells could continue to support phage replication and later increase in frequency through growth or conjugative transfer. Thus, after the acute competition between lysis and host proliferation, longer-term viral pressure may reshape the frequency of plasmid states within the population. This feedback could generate negative frequency-dependent selection, maintaining dynamic variation in plasmid carriage and viral load without fixation of chromosomal resistance mutations.

Beyond this experimental system, such dynamics may help explain variable plasmid carriage in clinical and environmental bacterial isolates. Rather than reflecting plasmid instability alone, differences in plasmid prevalence may represent snapshots of dynamic cycles shaped by phage predation, plasmid costs, and horizontal transfer [79,80]. More broadly, our results suggest that bacterial survival during RNA phage infection can depend not only on preventing infection, but also on altering the timing of cell death and the population structure on which phage selection acts.

## Methods

### MS2 bacteriophage stock preparation

MS2 phage (ATCC 15597-B1) was propagated on log-phase *E. coli* C-3000 donor cells (ATCC 15597) in LB medium at 37°C with shaking overnight. Cells were pelleted by centrifugation at 4000 rpm for 20 minutes at 25°C. The supernatant was then filtered through a 0.22-μm Stericup filter (EMD Millipore) to remove any remaining cellular debris. The purified phage stock was stored at 4°C, typically achieving a concentration of 10^11^ to 10^12^ PFU/mL.

### Growth and infection conditions

*E. coli* was cultured in LB medium at 37°C with shaking. Once the culture reached an optical density at 600 nm (*OD*_600_) of 0.5, samples were either infected with MS2 bacteriophage or treated with LB medium for mock infection. Based on a calibration performed in our laboratory, an *OD*_600_ of 0.5 corresponds to approximately 10^8^ cells/mL; the volume of phage stock required for each MOI was calculated accordingly, using the stock titer determined by plaque assay. For infected samples, MS2 bacteriophage was added at a multiplicity of infection (MOI) of 10 unless otherwise specified. Cultures were maintained at 37°C with shaking at 200 rpm. For total RNA extraction or scRNAseq processing, 1 mL of *E. coli* cells was collected from each culture and processed downstream as described below.

### Plaque Assay

The plaque assay was used to determine the concentration of the MS2 phage (virus titer). *E. coli* was cultured to mid-logarithmic phase (*OD*_600_ of 0.5) in LB medium at 37°C with shaking. Serial dilutions of the MS2 samples were prepared in 0.85% NaCl to reduce the phage concentration to below 200 PFU/ 0.1mL, ensuring plaques could be easily counted on each plate with the naked eye. Approximately 5 mL of soft agar was dispensed into each test tube, along with 1 mL of *E. coli*. Subsequently, 0.1 mL of each diluted MS2 phage sample was added to the tubes. The mixtures were poured onto petri dishes containing solid base agar and left to harden. The plates were then incubated overnight at 37°C and plaques were counted to calculate MS2 phage concentration in PFU/mL.

### Counting colony-forming unit (CFU)/mL

*E. coli* was cultured to mid-logarithmic phase (*OD*_600_ of 0.5) in LB medium at 37°C with shaking. A 1 mL sample of the culture was pelleted by centrifugation at 13,000 rpm for 1 minute at 4°C. The pellet was resuspended in 1 mL of LB medium and then serially diluted to achieve the appropriate concentrations. A 0.1 mL volume of each dilution was spread onto solid LB agar plates and left to harden. The plates were then incubated overnight at 37°, and CFUs were manually counted. The CFU/mL was calculated using the formula: 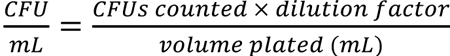. The estimation of surviving cells at 120 minutes post-infection was calculated as the ratio of infected to control CFU at this 120 minute timepoint. This 2% ratio serves as an estimate of the surviving fraction while accounting for host replication.

### Real time PCR

For cDNA synthesis, 500 ng of purified RNA was reverse-transcribed using SuperScript IV Reverse Transcriptase following the manufacturer’s protocol. Primers (2F-forward) GGCACAAGTTGCAGGATGCA and (2R-reverse) CGTGTCTGATCCACGGC targeting the MS2 genome were used to reverse-transcribe the positive and negative strands viral RNAs, respectively. Reverse transcription was also performed using random hexamers for the 16S quantification, used to control for the number of cells. qPCR was performed using iTaq Universal SYBR Green Supermix in Bio-Rad CFX Real-Time PCR. MS2 primers (2F and 2R) were used for forward and reverse MS2 quantification, and primers targeting the 16S gene (forward: AGCTAACGCGTTAAGTCG, reverse: TAAGGTTCTTCGCGTTGC) were used for 16S quantification. Each reaction contained 0.5µl of the cDNA along with gene-specific primers (final concentration of 500nM each). Samples were run in technical triplicates using the manufacturer’s protocol.

### Bacterial growth curves

Bacteria were grown to an optical density at 600 nm (*OD*_600_) of 0.5 and then divided to microplates at 37°C in LB medium. Wells in microplates were immediately infected by either MS2 or LB for mock infection. Conditions were kept at 37°C and *OD*_600_ was measured every 15 minutes over a 12-hour period using a BioTek plate reader. Plates were shaken just before every measurement. Each condition was measured in three technical replicates. Blank LB measurements were subtracted from bacterial OD measurements before analysis.

### Total RNA extraction

Bacteria isolated from experiments were immediately centrifuged at 13,000 rpm for 1 minute at 4°C, supernatant was removed and the pellet saved in −80℃ for downstream processing. Total RNA from *E. coli* cells containing both host and bacteriophage RNA was extracted using the RNeasy Mini Kit (Qiagen) following the manufacturer’s protocol for bacterial RNA purification (“Purification of Total RNA from Bacteria Using the RNeasy® Mini Kit - (EN)”). The initial bead-based lysis step was modified to a chemical lysis using 20 mg/mL of lysozyme (catalog number: Sigma-Aldrich L6876), prepared in TE buffer. After adding the lysozyme solution, samples were vortexed for 10 seconds every two minutes, over a total duration of 10 minutes.

### Sample preparation and sequencing of bulk RNA-seq

500 ng of total RNA was used for rRNA depletion and directional mRNA sequencing using NEBNext® rRNA Depletion Kit (Bacteria) followed by NEBNext® Ultra™ II Directional RNA Library Prep Kit for Illumina. Illumina adaptors for sequencing were added using NEBNext® Multiplex Oligos for Illumina® kit, and samples were sequenced on Illumina NextSeq platform using either 1×75bp for the main time-course experiment, and 1×100bp for the isolate RNA-seq experiment, aiming for 10 million reads per sample.

### Bulk RNA-seq computational analysis

First, adaptor contamination was removed from reads and low quality bases were trimmed using Trimmomatic [81] with the following parameters: “ILLUMINACLIP:TruSeq3-SE.fa:2:30:10 SLIDINGWINDOW:5:15” (with an additional “MINLEN:36” parameter applied specifically to the isolate RNA-seq experiment).

Next, the reads were aligned using Burrows-Wheeler Aligner (BWA) [82] to the C-3000 and MS2 genomes (FASTA sequence and annotations downloaded from ATCC Genome Portal [83]). For C-3000 annotations, alignment to the Swiss-prot database [84] was used to supplement annotations of putative proteins in the ATCC annotation. Unannotated, putative proteins remained denoted as LELOBEKK_x. Reads were filtered using SAMtools [85] to exclude unmapped reads and retain alignments with a MAPQ score of 30 or higher. Reads aligned to sequences annotated as coding (CDS) were then counted using Htseq-count [86] (parameters: stranded=reverse, idattr=Name, type=CDS). Because the libraries were strand-specific, reads mapping to the MS2 genome were additionally quantified according to strand orientation to obtain separate positive- and negative-strand viral RNA counts. After creating a counts matrix per experiment, samples were analyzed using the standard DESeq2 [87] pipeline. Counts were normalized using the DESeq2 median-of-ratios method, and a global model was fitted with the design ∼ line + time_scaled + infected, where line denotes the biological replicate line, included as a blocking factor, and time_scaled is sampling time centered and scaled. Differential expression for specific contrasts was assessed by independent pairwise DESeq2 analyses, each fitted to the relevant subset of samples with size factors and dispersions estimated within that subset: infected versus control at each matched time point (design ∼ line + infected), and, within each infection condition separately, 45 versus 15 minutes and 180 versus 15 minutes post-infection (design ∼ line + time). Genes were called differentially expressed at a Benjamini-Hochberg adjusted P value below 0.05 together with |log2 fold change| > 1.5; MS2 features were excluded from these counts. For the 180-versus-15 comparison shown in Fig. S3, genes that changed in the same direction in control cultures over the same interval were further subtracted, so that the reported counts reflect changes specific to infection rather than shared culture-density effects; the 45-versus-15 comparison is instead shown separately for infected and control cultures. A regularized log transformation was applied to the count data for sample clustering and visualization.

Size factors were estimated with MS2 features retained in the count matrix. This has no effect on normalization: the median-of-ratios method uses only genes with a non-zero count in every sample, and both viral features had zero counts in multiple control samples, so they were excluded from the size-factor calculation by construction. Recomputing size factors after removing the MS2 rows returned identical values.

For the isolates RNA-seq experiment, transcriptomic variation among phage-exposed and control colony-derived populations was assessed using normalized expression data, including MS2 and F-plasmid transcripts.

Of note, we observed very minor MS2 contamination in the time-course control samples, at most 0.01% of mapped reads in any control sample, approximately four orders of magnitude below the proportion in time-matched infected samples (Fig. S14). In the most conservative view, such contamination would be expected to reduce, rather than inflate, differences between infected and control samples. Additionally, minor MS2 presence was observed in the colony bulk RNA-seq data, where MS2 reads accounted for no more than 0.03% of the total mapped reads in each of the three control samples.

### Functional enrichment for gene set

Functional enrichment was performed using the STRING database [88], with the *E. coli* K-12 gene set used as the reference background because C-3000 is derived from K-12. Gene Ontology Biological Process terms were prioritized; Cellular Component terms were used where no informative Biological Process enrichment was available.

### F-plasmid gene modules

F-plasmid genes were split into two modules. The *tra* module contained the genes of the F transfer region, which encode F-pilus biogenesis, mating-pair stabilization and DNA transfer (all *tra* and *trb* genes, together with *finO*; n = 33 genes). The non-tra module contained all other annotated F-plasmid coding genes (n = 53), including genes for replication, partitioning and plasmid maintenance. For the bulk time course, the module score of each sample was the mean normalized count (DESeq2 size-factor normalized) across the genes in that module. For the colony isolates, genes with fewer than 10 reads in at least three samples were excluded, leaving 79 expressed F-plasmid genes (33 *tra* and 46 non-tra), and the module score was the mean across member genes of log10(CPM + 1), where CPM is computed relative to host-mapped reads, with library size factors estimated from chromosomal genes only.

### Colony isolation for whole-genome sequencing and bulk RNA-seq

Host cultures (50 mL) were grown as described in the “Growth and infection conditions” section, and either challenged with 5 mL of MS2 at an MOI of 10 or mock infected with 5 mL of LB medium to serve as unexposed controls. Samples were then plated as described in the “Counting colony-forming unit (CFU)/mL” section. Following overnight incubation, 10 single colonies were randomly selected from the control plates, and 20 single colonies were selected from the phage exposed plates, with each colony used to establish a separate clonal isolate. Of the 20 phage exposed lines, half were derived from cultures plated immediately following infection, and the remaining half were plated 180 minutes post-infection. Because both subgroups underwent subsequent overnight incubation in the presence of MS2 on the agar surface, they were treated as a single experimental group; no significant differences in mutation profiles or plasmid coverage were observed between the two phage-exposed subgroups (not shown). Frozen stocks were established for each of the 30 isolates by mixing 500 µL of an overnight starter culture with 500 µL of sterile 30% glycerol. For all downstream experimental assays, frozen stocks were streaked onto fresh plates, and individual colonies were used to initiate cultures.

### Genomic DNA extraction

Genomic DNA was extracted from the 30 isolated *E. coli* colonies (20 phage exposed isolates and 10 unexposed controls) derived from overnight starter cultures. Total genomic DNA, encompassing both chromosomal and plasmid DNA, was isolated using the DNeasy UltraClean Microbial Kit (Qiagen) according to the manufacturer’s protocol for bacterial genomic DNA purification.

### Whole-genome sequencing

Bacterial Genome Sequencing was performed by Plasmidsaurus using Oxford Nanopore Technology (ONT). Raw FASTQ files were generated and used for all downstream analysis. Long read sequencing data in FASTQ format were aligned to the *E. coli* C-3000 (ATCC 15597) reference genome using minimap2 (version 2.31) [89] utilizing the Oxford Nanopore preset (-ax map-ont). Secondary alignments were discarded (-- secondary=no), and soft clipping was applied for supplementary alignments (-Y). The resulting alignments were immediately converted, sorted, and indexed using SAMtools (version 1.23.1) [85] to generate binary alignment map (BAM) files. Single-nucleotide variants (SNVs) and short insertions/deletions (indels) were identified from the BAM files using LoFreq (version 2.1.5) [90]. Prior to variant calling, uniform indel quality scores of 16 were assigned using the LoFreq indelqual module. Raw variant calling was executed with default filtering disabled and indel calling enabled. The resulting calls were subjected to initial statistical filtering via LoFreq filter, retaining variants that met a minimum read coverage of 30x, a minimum allele frequency of 0.01 (1%), and passed a strand-bias p-value threshold of 0.01. For analyses of fixed genetic changes, variants were subsequently restricted to those reaching > 80% within a colony-derived population.

To eliminate false-positive indel calls in low-complexity genomic regions, a custom Python (version 3.10.20) filtering step was applied. For each indel, the Shannon entropy of the adjacent reference sequences flanking the mutation site was calculated. Indels with the product of the forward and reverse entropies score below 1.0 were classified as sequencing artifacts and filtered out, whereas all SNVs automatically bypassed this filter. Concurrently, structural variants (SVs) were called from the sorted BAM files using Sniffles (version 2.6.2) [91] against the reference genome using default parameters. To ensure variant call robustness, samples were additionally evaluated using Medaka by ONT Research (version 2.0.1; https://github.com/nanoporetech/medaka), breseq (version 0.39.0) [92], and bcftools (version 1.22) [85]; because these secondary pipelines yielded highly concordant mutation profiles, differing only slightly in estimated mutation frequencies according to each algorithm, these validation datasets are omitted. The orientation of the *fimS* invertible promoter element was additionally examined from the long-read alignments to assess phase variation of type I fimbrial expression. All final variants were compiled into standard Variant Call Format (VCF) file for downstream comparative analysis.

For plasmid copy-number estimation, reads were aligned with minimap2 (version 2.30, -ax map-ont), and mean per-base depth of the chromosome and the F-plasmid was computed from the sorted alignments using pysam (https://github.com/pysam-developers/pysam).

### Plasmid to chromosome copy number estimation

To estimate relative plasmid to chromosome copy number ratios from the whole-genome sequencing reads, binary alignment map (BAM) files were analyzed using a custom Python script. Sequencing depth was calculated on a per-base level across both the host chromosome and F-plasmid reference. Relative plasmid abundance was determined by normalizing the mean coverage depth of the F-plasmid to that of the host chromosome.

### scRNAseq sample preparation and sequencing

Bacteria isolated from experiments were immediately fixed using 1% formaldehyde. Cells were centrifuged, and pellets were resuspended in 1 mL MAAM (4:1 V:V dilution of methanol to glacial acetic acid). Samples were kept at −20℃ until probe hybridization. Downstream processing and sequencing were performed as described in the protocol [36]. The study comprised two main experiments. In the first, time points 15, 30, and 45 minutes were sequenced with associated controls; libraries were sequenced on the Illumina platform for 28 bases in the R1 direction and 151 bases in the R2 direction. In the second, time points 15, 45, 120 and 180 minutes were sequenced with associated controls; in this experiment, MS2 probes were read out as a separate feature-barcode library, which allowed more of the sequencing depth to be allocated to host mRNA while retaining viral counts for each cell, and libraries were sequenced on an Element AVITI for 28 bases in the R1 direction and 150 bases in the R2 direction. However, sequencing failed for the infected 15 and 45 minute samples and the 120 and 180 minute controls of the second experiment. Consequently, we used only the 120 and 180 minute samples to estimate MS2 abundance (which was very high across all cells); given their comparable viral read fractions, only the 180 minute time point is presented here.

### Probe design for scRNAseq

Approximately 50 nucleotide long probes were designed with a region complementary to the mRNAs of the *E. coli* genome and to the MS2 genome. The probes also included a poly-A tail for capture on the 10x Genomics platform and a unique molecular identifier (UMI) for removal of PCR duplicates. To improve transcript capture efficiency and minimize noise from inadequate hybridization or weak amplification of individual probes, multiple probes targeting different regions of each gene were designed. The probe pool [93] used was originally designed for the closely related *E. coli* MG1655 (21,527 probes for 4181 genes) and supplemented with probes complementary to the MS2 genome for both the positive and the negative strands (26 and 25 probes, respectively). Probes for the 50 most differentially expressed C-3000 genes from the bulk RNA-seq results (which were absent in the original MG1655 pool) and probes targeting the F-plasmid encoded *tra* operon were also added to the pool (Table S7). In the second round of scRNAseq, some of the probes were augmented and corrected (Table S8).

### scRNAseq read preprocessing and alignment

In both experiments, the 10x bead-derived UMI was replaced with the probe-encoded UMI, because PCR within droplets can generate new bead-derived UMIs, whereas the probe-encoded UMI is carried by the original hybridized probe.

#### First experiment

Samples were analyzed as described in the ProBac-seq toolkit (https://gitlab.com/hormozlab/probacseq-toolkit/-/blob/main/example.ipynb). Briefly, the 10x bead UMI was replaced with the probe-encoded UMI, and reads containing the cell barcode, the UMI and the probe sequence were mapped using the standard 10x Genomics Cell Ranger pipeline [94].

#### Second experiment (180 minute sample)

In this library design, R1 (28 bp) contains the 16-bp 10x cell barcode followed by a 12-bp bead-derived UMI, and R2 contains the hybridized probe flanked by two constant sequences (5′ anchor GATGACCCGGTCCATAC and 3′ anchor CATAGTTTCTTCGAGCAAGCTT), followed by the probe-encoded UMI and the poly(A) tail. Using splitcode (v0.30.0) [95], a modified R1 was generated by joining the first 16 bases of R1 (cell barcode) with the 12 bases immediately downstream of the 3′ anchor in R2 (up to 3 mismatches, searched within R2 positions 50–110). Read pairs in which the anchor was not detected (modified R1 shorter than 26 bases) were removed with Trimmomatic (v0.39) in paired-end mode (MINLEN:26). R2 was then trimmed to the probe sequence located between the 5′ anchor (up to 1 mismatch, positions 0–50) and the 3′ anchor (up to 3 mismatches, positions 50–150) using splitcode. For the gene-expression library, trimmed R2 reads were reverse-complemented (Biopython) to match the transcript orientation expected by Cell Ranger; the MS2 feature-barcode library was retained in probe orientation. Gene-expression and MS2 feature-barcode libraries were analyzed jointly with Cell Ranger multi (v9.0.1) [94] using a custom *E. coli* C-3000 probe reference (excluding MS2) built with CellRanger mkref, with MS2 probes quantified as Custom feature barcodes defined by their probe sequences.

### scRNAseq computational analysis

Matrices contained counts of every probe per cell. Analysis used the raw CellRanger matrices rather than the filtered ones, since CellRanger’s cell calling is tuned for eukaryotic cells with thousands of UMIs; barcodes with fewer than three total UMIs were discarded at load. Probes were collapsed to gene level by taking, for each gene and cell, the maximum count across the probes targeting that gene. Quality metrics were computed on the gene-level matrix, using host (non-MS2) features only, so that infected cells with reduced host transcription were not preferentially discarded. Barcodes were retained if they had at least 50 detected host genes and a host count total of at least 100. Because genes are summarised by the maximum across their probes, this total is a sum of per-gene maxima rather than a UMI count; it serves only as the filtering statistic, while every ratio and normalisation offset below uses the probe-level host UMI total. If a library’s 1st percentile exceeded either threshold, the library-specific percentile was used instead; barcodes were also required to fall below the 99th percentile for both metrics. Genes detected in fewer than three cells were removed, as were a predefined list of 18 outlier genes (Fig. S5); MS2 features were retained regardless of both criteria, so that the feature set is identical across samples even where MS2 is near-absent. Counts were normalized using host (non-MS2) features only. Each gene-level value was scaled by the ratio of the median host total across cells to that cell’s own host total, then log-transformed; the host total here is the sum of a cell’s gene-level values.

The 1,000 most variable host genes were used for PCA (20 components computed, the first 10 retained), shared-nearest-neighbour graph construction (k = 20), Louvain clustering (resolution 0.2) and UMAP (30 neighbours). MS2 features were excluded from variable-gene selection so that the embedding reflects host state rather than viral load. Each sample was processed independently, so cluster identities are not comparable across samples. Cluster markers were identified with a Wilcoxon rank-sum test of each cluster against all others. Analysis was performed in R (v4.3.3) using Seurat v5.3.0 [96].

To classify infection states in the first experiment, background viral signal was modelled from the percentage of MS2 UMIs relative to the sum of MS2 and host UMIs, both taken from the probe-level matrix, in the uninfected control cells, pooled across timepoints. Cells above the 95th percentile of this control distribution were classified as ambiguous, and cells exceeding the 99th percentile were classified as infected provided they also met a structural validation criterion: at least seven distinct MS2 probes, drawn from the 26-probe subset targeting the positive strand, detected in the same cell. Cells meeting the percentile criterion but not the probe criterion remained ambiguous. Cells at or below the 95th percentile were classified as uninfected. For all downstream analyses, ambiguous and uninfected cells within infected libraries were grouped together and are referred to as exposed-uninfected, i.e. cells that were exposed to phage but showed no confident evidence of infection.

The 180 minute sample of the second experiment was processed without matched controls, so no contamination null was available, and the threshold was derived from the shape of the MS2 distribution itself, using Otsu’s method on the log10 ratio of MS2 to host UMIs across cells with at least one MS2 UMI; for this sample, MS2 UMIs were taken from the MS2 feature-barcode library. The ratio was used in place of percent MS2 because the latter saturates once phage dominates the transcriptome.

For pseudo-bulk differential-expression analysis, cells were grouped by biological sample and infection-state classification, and each group was split at random into four pools. Probe counts were summed within each pool and across the probes of each gene. Differential expression between groups was performed with DESeq2 [87] under a ∼group design, with size factors set to each pool’s host UMI total normalised to its geometric mean. Each module’s effect was then calibrated against 2,000 random gene sets drawn from the same probe panel, matched in size and stratified into ten expression-rank bins so that the comparison genes were of comparable abundance; the reported significance is the position of the observed effect within that null.

Expression of the F-plasmid transfer region was compared within each infected library, between cells called infected and exposed-uninfected cells. The module comprises the 15 transfer-region genes represented on the probe panel. The module total and each gene individually were tested with a negative-binomial GLM using log host UMIs as offset, and per-gene p-values were Benjamini–Hochberg corrected within each timepoint. Per-gene results are shown for the 13 genes reaching at least 150 UMIs at one or more timepoints.

### Single-molecule fluorescence in situ hybridization (smFISH)

smFISH detection of intracellular MS2 RNA was performed based on the protocol previously described [50]. Briefly, a set of oligonucleotide probes complementary to the positive-sense MS2 genome was used to detect viral RNA within individual *E. coli* cells. At the indicated time points following infection, cells were collected and fixed in formaldehyde, washed in phosphate-buffered saline, and permeabilized with ethanol.

Cells were subsequently incubated in formamide-containing wash buffer and hybridized overnight with the MS2 probe mixture in hybridization buffer. Following hybridization, cells were washed to remove unbound probe, stained with DAPI, and resuspended for microscopy. Imaging was performed using a Nikon Eclipse Ti2 inverted epifluorescence microscope equipped with a 100× oil-immersion objective (Plan Apochromat, NA 1.40) and a Hamamatsu Flash4.0 camera (C11440-22CU). Phase-contrast and fluorescence images were acquired. Quasar 570-labeled smFISH probes were imaged in the TRITC channel, and DAPI was imaged in the DAPI channel for DNA visualization. Z-stacks were acquired at 0.2-µm intervals using identical acquisition settings across all samples.

Additional microscopy fields were subsequently acquired from the same fixed samples to increase cell numbers at sparsely sampled time points. These images were analyzed separately, using an MS2-positive threshold determined from negative-control images acquired in the same imaging session (Table S3).

### smFISH computational analysis

For each field, the best-focus plane was manually selected from the phase-contrast Z-stack based on cell morphology and used for cell segmentation. Individual cells were segmented using Cellpose-SAM (version 4.1.1) [97] with a cell diameter of 22 pixels, a flow threshold of 0.4 and a cell-probability threshold of 0, and the resulting cell masks were used for whole-cell fluorescence quantification. During initial calibration of the segmentation, the DAPI signal was used to manually verify that the segmented masks corresponded to cells. Cells touching the image border were excluded. Because the MS2 signal at later time points was too intense for spot-based quantification [40], we adapted this approach to use whole-cell fluorescence. MS2 fluorescence was quantified from the mean projection of the TRITC Z-stack. For each cell, the integrated signal within the mask was background-corrected by subtracting the median intensity of a ring 0.2–0.5 µm around the cell, excluding pixels belonging to any segmented cell, multiplied by the cell area. The corrected signal was normalized to cell area (in pixels) to obtain the whole-cell signal density. Cells were classified as MS2-positive if their signal density exceeded the maximum observed in negative-control cells (Fig. S10). The percentage of MS2-positive cells was calculated for each microscopy field and across pooled cells for each time point, with 95% confidence intervals for pooled percentages computed using the Wilson score method.

For representative images, the best-focus plane was selected from each Z-stack. Identical linear display settings were applied to the time-course images, whereas display ranges for images of dividing cells were adjusted individually for visualization.

### Deletion strains

Strain construction was achieved via conjugation between a recipient strain from the KEIO [59] (knockout) *E. coli* BW25113 collection and an F′ donor strain, XL1-Blue. The XL1-Blue strain carries tetracycline resistance, and the KEIO strains are kanamycin resistant. Overnight cultures of both donor and recipient strains with knockout of genes of interest were diluted in LB medium, and 200 µl of log-phase donor cells were combined with 200 µl of log-phase recipient cells (*OD*_600_ = 0.4). The mixture was incubated at 37°C for two hours without shaking. Afterward, samples were vortexed to halt conjugal transfer, and bacteria were plated on LB agar containing both tetracycline and kanamycin to isolate transconjugant cells that acquired the F-plasmid. Plates were incubated overnight, and a single colony was subsequently isolated and used for cell density experiments.

### Endpoint phage titer assay

Overnight starter cultures of the knockout strains were measured for optical density at 600 nm (*OD*_600_) and diluted into 20 mL of fresh LB medium to establish uniform initial cell densities. Cultures were grown at 37°C with shaking until reaching *OD*_600_ = 0.5. For infection assays, 5 mL aliquots of each strain were transferred into 50 mL tubes and infected in triplicate with MS2 bacteriophage at MOI of 0.1 and 10. Infected cultures were then incubated at 37°C for 3 hours.

Following incubation, 1 mL was sampled from each culture and centrifuged to pellet bacterial cells. The phage containing supernatant was collected, serially diluted tenfold in LB medium, and viral titers (PFU/mL) were quantified via plaque assay as described above. The relative endpoint titer was defined as the PFU titer recovered from each deletion strain after 3 hours of infection divided by the PFU titer recovered from the control strain (*ΔlacY*), under the same MOI conditions.

## Data and Code Availability

The source code is available at https://github.com/Stern-Lab/MS2_C3000_recovery. Sequencing data have been deposited in GEO under SuperSeries accession GSE349161 and in the SRA under BioProject accession PRJNA1536727. The *E. coli* C-3000 and MS2 reference genomes are available from the ATCC Genome Portal (https://genomes.atcc.org). Supplementary Tables S1, S5, S7 and S8 are available at Zenodo (https://doi.org/10.5281/zenodo.23232244).

## Acknowledgements

We thank Avigdor Eldar and Omri Wurtzel for critical discussions; and Lanying Zeng for the kind gift of smFISH probes for MS2. This study was supported by an ERC starting grant 852223 (RNAVirFitness) to A.S. IG was supported by the National Institutes of Health grant R35 GM140709, the National Science Foundation grant 2243257 (NSF Science and Technology Center for Quantitative Cell Biology), and the Alfred P. Sloan Foundation grant G-2023-19649. This study was also supported by a fellowship to N.H. and Y.R. from the Edmond J. Safra Center for Bioinformatics at Tel Aviv University.

## Notes

https://www.ncbi.nlm.nih.gov/geo/query/acc.cgi?acc=GSE349161

https://www.ncbi.nlm.nih.gov/sra/?term=PRJNA1536727

https://doi.org/10.5281/zenodo.23232244

https://github.com/Stern-Lab/MS2_C3000_recovery

### Competing Interest Statement

The authors have declared no competing interest.

