## Supplementary Material for "Temporal dynamics of RNA phage infection reveal bacterial recovery despite ongoing viral replication"

### Supplementary Material for Harel et al.

#### Supplementary figures

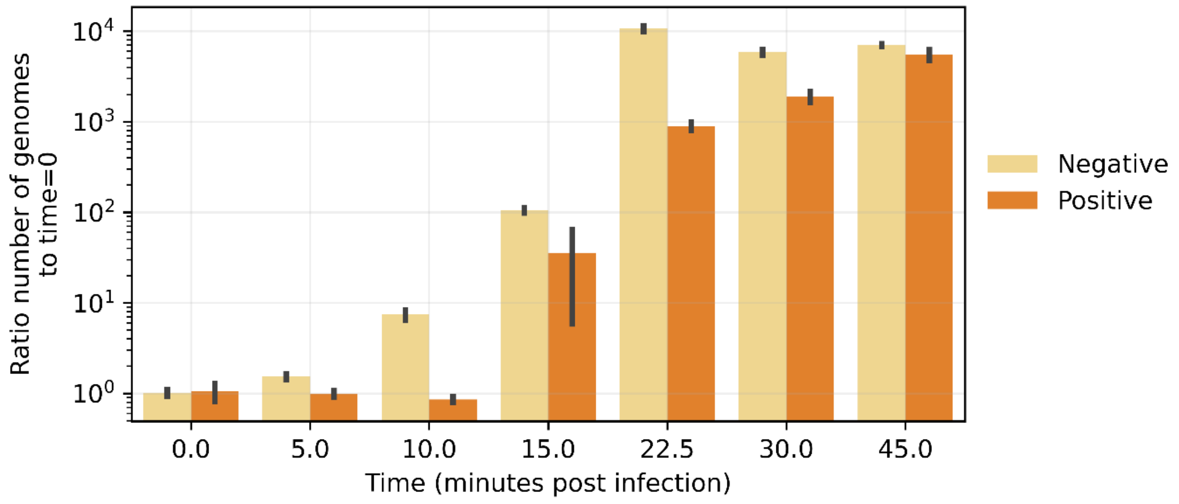


**Figure S1. Early kinetics of MS2 RNA strand synthesis.** Intracellular accumulation of MS2 negative (beige) and positive (orange) RNA strands quantified by real-time PCR across the first 45 minutes post-infection. Cq values were normalized to 16S rRNA in the same sample and expressed as fold change relative to time point 0, shown on a logarithmic scale. Error bars denote the standard error of the mean of technical replicates. Negative-strand RNA rose first, increasing by ~10-fold within 10 minutes and peaking at ~22.5 minutes, at which point its fold change exceeded that of the positive strand by over an order of magnitude. Positive-strand RNA accumulated later, expanding steeply between 15 and 45 minutes until the two strands reached comparable fold changes by 45 minutes. These dynamics indicate that negative-strand synthesis precedes and is largely completed before the bulk of positive-strand production, consistent with the stamping-machine mode of replication (Fig. 1B, C).

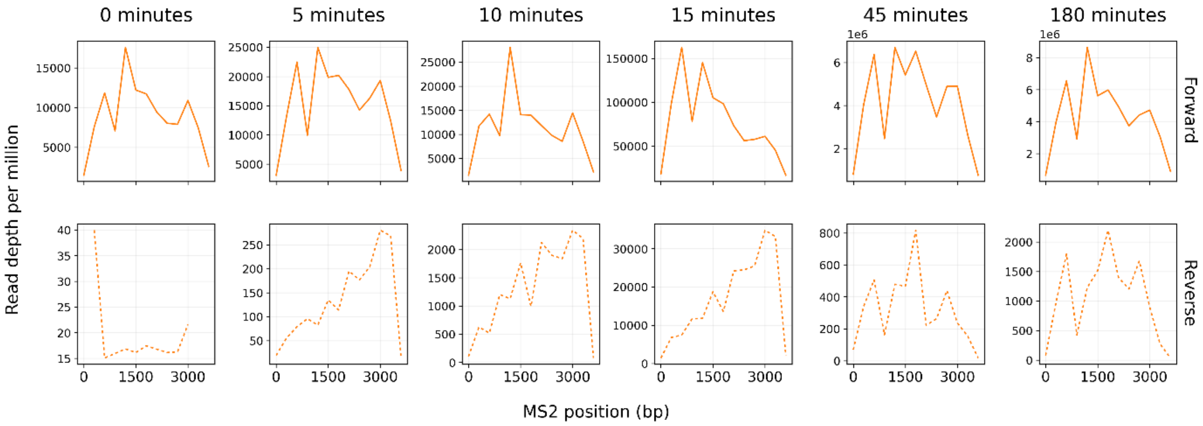


**Figure S2. Strand-specific coverage across the MS2 genome during infection.** Coverage of positive- and negative-strand MS2 RNA is shown across the viral genome at the indicated time points. Negative-strand RNA displays pronounced enrichment toward the genomic 3′ end during early infection. Because negative-strand synthesis initiates at the 3′ end of the positive-strand genome, this pattern is compatible with an abundance of nascent, incompletely synthesized negative strands and therefore with ongoing negative-strand synthesis. However, preferential degradation or differential stability along negative-strand RNA could produce a similar coverage bias, and the pattern therefore cannot uniquely distinguish synthesis from degradation. By later time points, negative-strand abundance declines while positive-strand RNA becomes highly abundant across the genome.

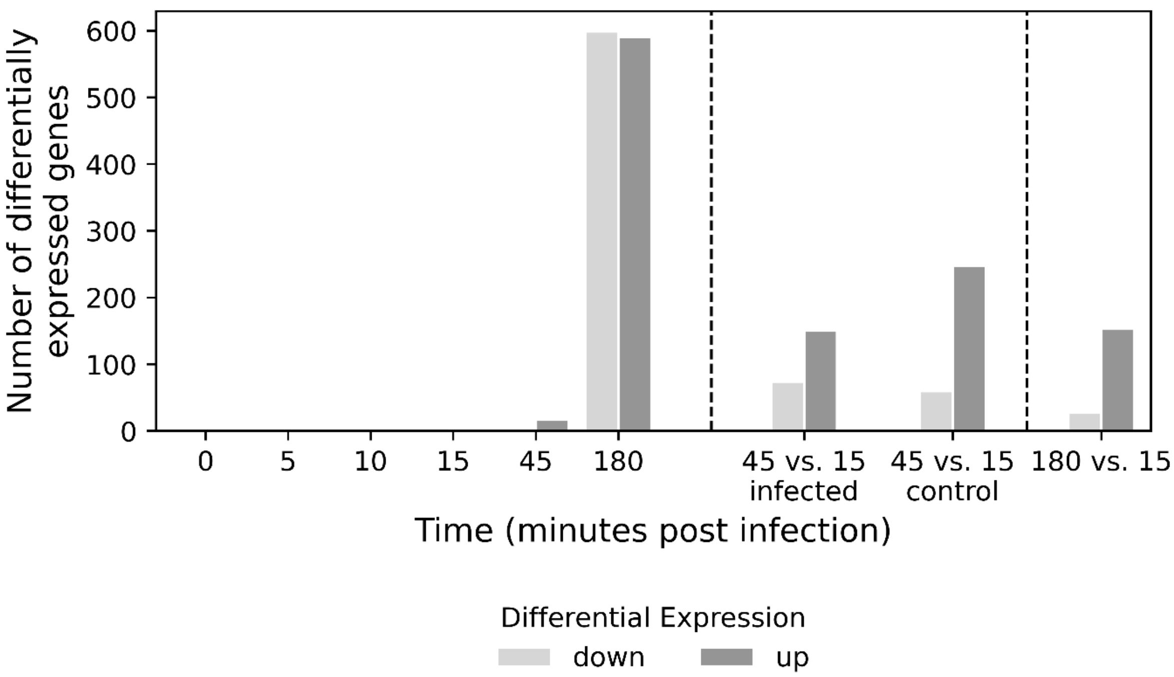


**Figure S3.** Counts of differentially expressed genes per time point, comparing the MS2-infected population to the control population. Genes were considered significantly up- or downregulated if $|log2\ fold\ change| > 1.5$ and adjusted p-value < 0.05. To the right of the first dashed line, temporal comparisons are shown: 45-minute samples compared with 15-minute samples, separately for infected and control populations, and 180-minute infected samples compared with 15-minute infected samples, excluding genes that changed in the same direction in controls over the same interval.

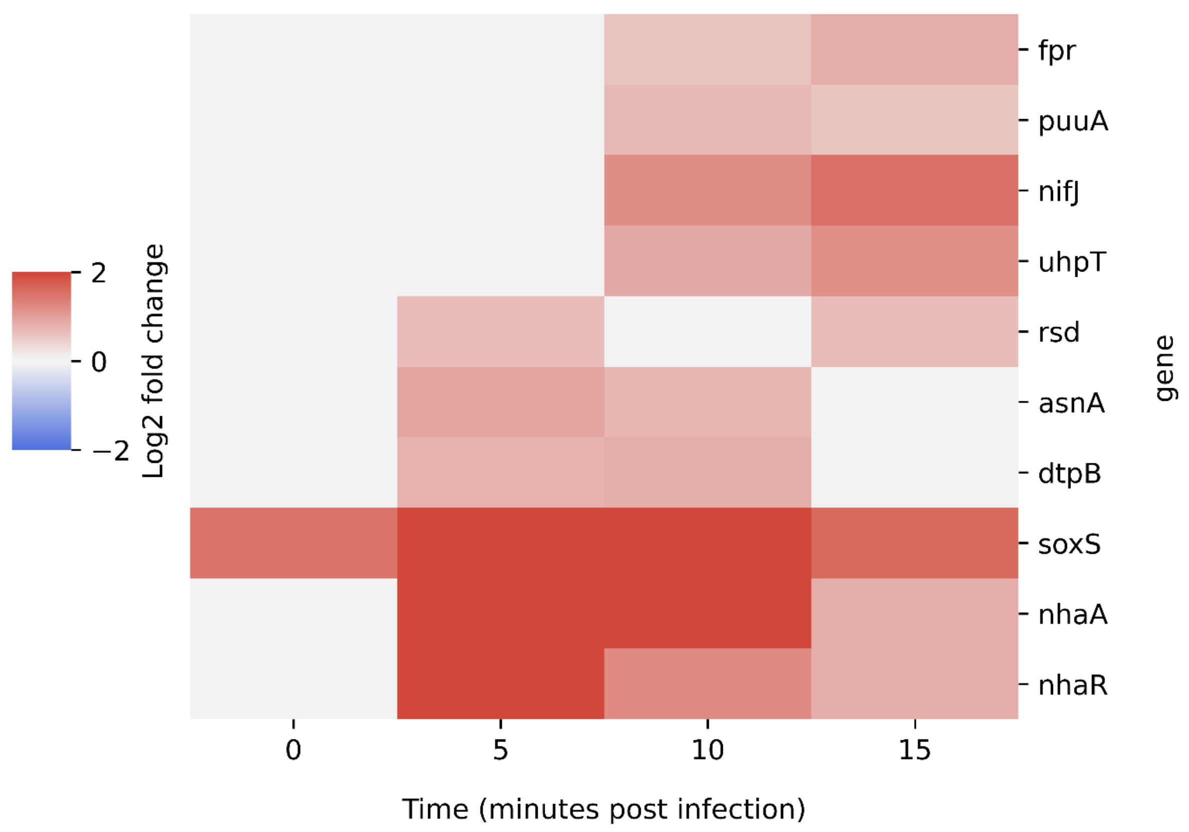


**nhaA** Na+:H+ antiporter

**nhaR** DNA-binding transcriptional activator

**rsd** regulator of sigma D (regulated by nhaR)

**puuA** glutamate-putrescine ligase

**asnA** asparagine synthetase A

**dtpB** dipeptide/tripeptide:H+ symporter (regulated by ppGpp)

**nifJ** pyruvate:ferredoxin (flavodoxin) oxidoreductase

**uhpT** hexose-6-phosphate:phosphate antiporter

**soxS** superoxide response transcriptional regulator SoxS

**fpr** flavodoxin/ferredoxin-NADP+ reductase (regulated by soxS)

**Figure S4. Genes upregulated at multiple early time points.** Upregulation of genes in earlier time-points of infection, shown are genes upregulated in more than one time point (0-15 minutes). Total of 10 genes. The strongest and most sustained induction was observed for nhaR and its target nhaA, which were elevated from 5 minutes onward. soxS was also strongly induced, but showed elevated expression already at time 0, before infection could elicit a transcriptional response, and was therefore not considered infection-specific.

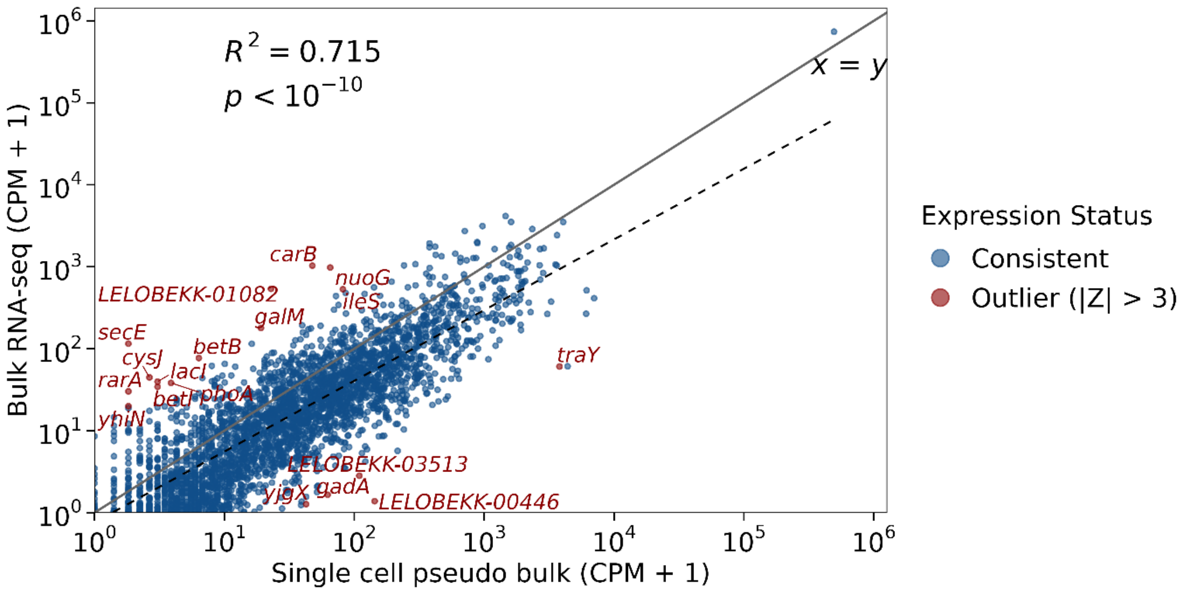


**Figure S5. Comparison of gene expression between single-cell pseudo bulk and bulk RNA-seq at 45 minutes post-infection.** Scatter plot showing normalized gene expression (counts per million (CPM) + 1) measured by single-cell pseudo bulk (x-axis) and bulk RNA-seq (y-axis) on $\log_{10}$ scales; each point represents a gene. Agreement between the two measurements was assessed by Pearson correlation of the $\log_{10}$(CPM + 1) values (r = 0.85, $R^2$ = 0.715, $P < 10^{-10}$). The solid gray line indicates the identity line (x = y) and the dashed black line the ordinary least-squares regression fit. Genes with standardized residuals |Z| > 3 are highlighted in red and labeled as expression outliers, whereas all other genes are shown in blue. Outlier classification was restricted to host genes, and the 18 identified host outliers were subsequently removed from downstream single-cell analysis.

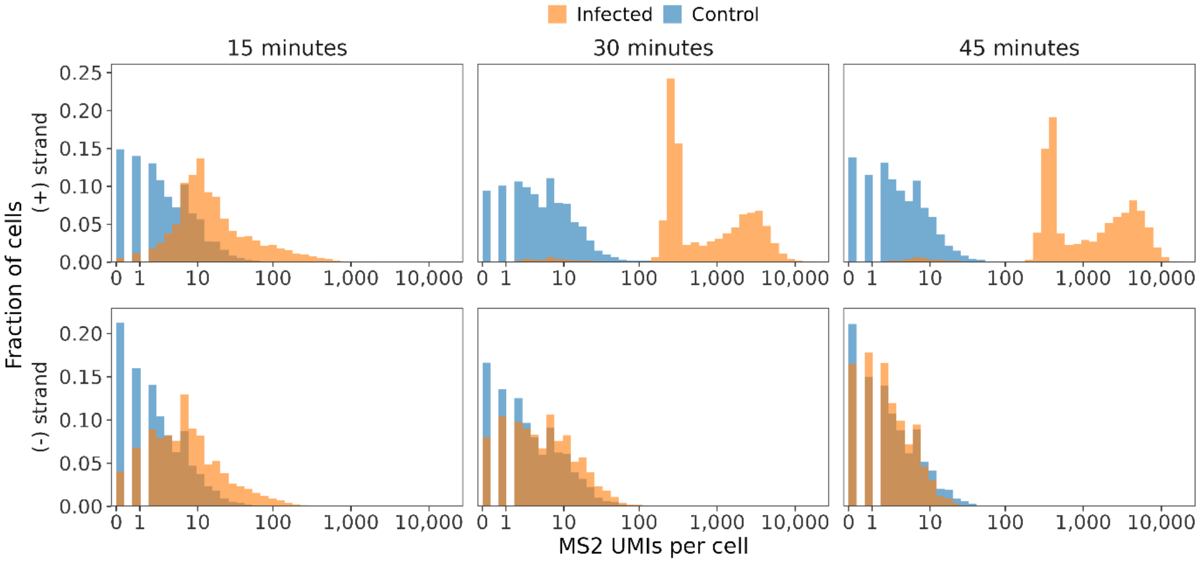


**Figure S6. Strand-specific MS2 signal separates infected from control cells only from 30 min, and only on the (+) strand.** Per-cell MS2 UMI counts for MS2-infected (orange) and control (blue) cells, shown as the fraction of cells against UMIs per cell on a log scale. Columns are timepoints (15, 30, 45 min); rows are the two probe sets: (+) strand and (−) strand. All host-filtered cells are included (n = 8,579 / 8,242 / 7,089 infected and 9,731 / 6,571 / 7,122 control at 15 / 30 / 45 min). On the (+) strand, infected and control distributions overlap at 15 min but resolve into two well-separated populations by 30 min and 45 min, where infected cells form a distinct high-copy mode (medians 371 and 893 UMIs/cell) far above the control (medians 5 and 4). The (−) strand shows no such separation at any timepoint (infected vs control medians 7/2, 5/3, 2/2 at 15/30/45 min), so it is uninformative for classifying infected cells at these timepoints. At 15 min, neither strand separates the two conditions, so infection status cannot be resolved reliably at the single-cell level at this early timepoint.

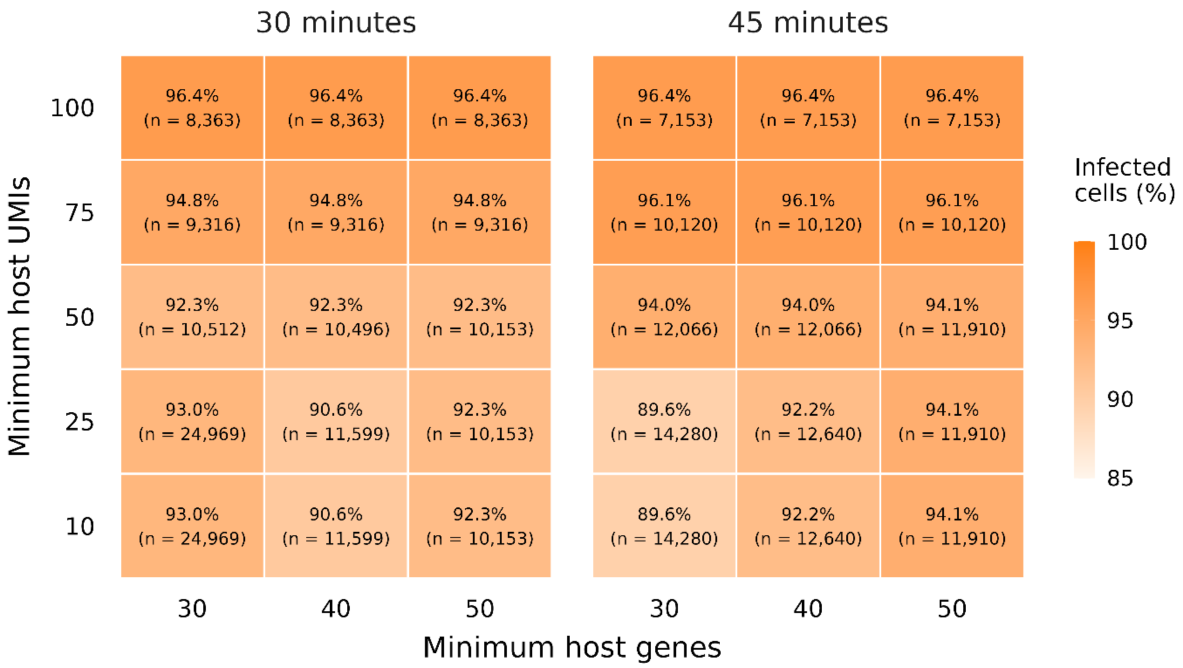


**Figure S7. The fraction of infected cells depends on the host-transcript QC thresholds.** Percentage of cells classified as MS2-infected in the 30- and 45-minute infected scRNA-seq libraries, across combinations of minimum host UMIs (rows) and minimum host genes (columns) per cell. Numbers in parentheses give the cells passing each setting. Depending on the thresholds, the infected fraction ranged from ~90% to ~96%, decreasing as more permissive cutoffs admitted cells with fewer host transcripts. The infection cutoffs were derived from the control libraries under the standard QC and kept fixed throughout.

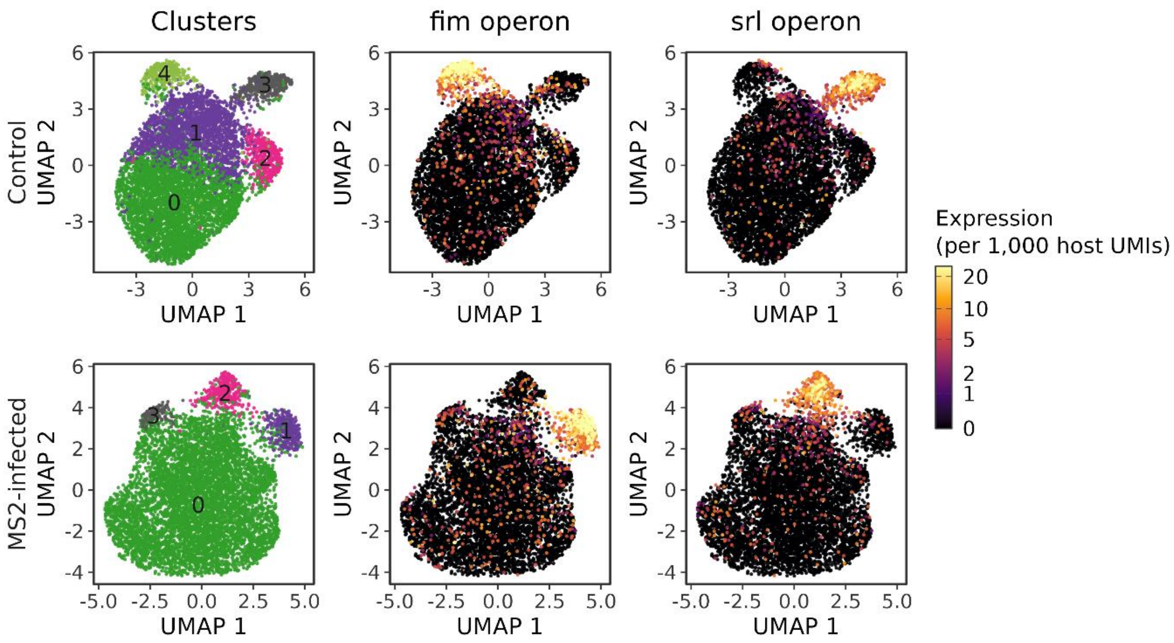


**Figure S8. UMAP visualization of scRNAseq data at 45 minutes.** Cells from the uninfected control (top) and the MS2-infected sample (bottom) are coloured by cluster identity (left), and by mean normalized expression of the *fim* (middle) and *srl* (right) operon genes, expressed per 1,000 host UMIs. Several distinct clusters are resolved in both samples, including a population with high *fim* operon expression (cluster 4 in the control, cluster 1 in the infected sample) and a population with high *srl* operon expression (cluster 3 in the control, cluster 2 in the infected sample). Because these populations are present in the control as well as in the infected sample, they reflect infection-independent heterogeneity, and we attribute the *fim*-high population to phase variation of the type I fimbrial switch; both operons were therefore excluded from downstream analyses. Cluster 3 of the infected sample corresponds to the exposed-uninfected population with low MS2 signal and elevated *mal*/*glp* expression shown in Fig. 2A.

(A)

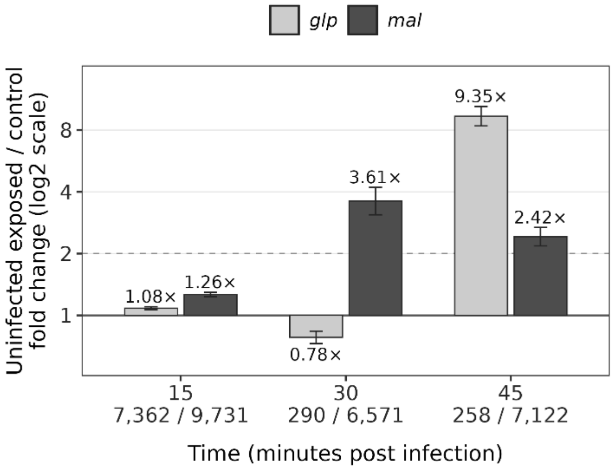


(B)

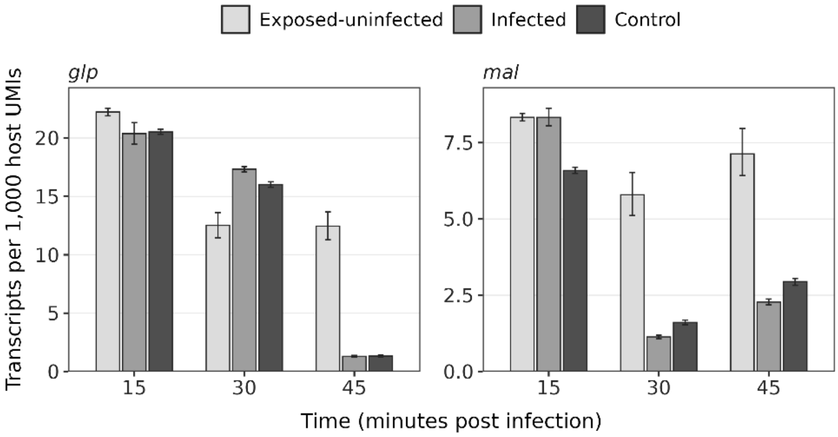


**Figure S9. *mal* and *glp* expression dynamics.** (A) Pseudo-bulk differential expression of the *mal* and *glp* regulons between exposed-uninfected cells and the time-matched uninfected control at 15, 30 and 45 minutes post-infection; numbers below each time point give cell counts (exposed / control). No significant difference was detected at 15 minutes; a subset of *mal* genes was elevated at 30 minutes, and both regulons were significantly higher at 45 minutes. (B) Summed *mal* and *glp* expression, normalized to host UMIs, in exposed-uninfected, infected and control cells (mean with bootstrap 95% CI). Expression declined sharply in infected and control cells but much less in exposed-uninfected cells, so the difference in (A) arises mainly from a lack of shutdown.

Two non-exclusive explanations could account for this signature. In the first, it is a bystander response to lysing neighbors: LB is a carbohydrate-limited medium dominated by amino acids, and lysis could release intracellular nutrients that neighboring cells take up through pathways such as *mal* and *glp*. In the second, it indicates carbon limitation: both regulons are catabolite-repressed and cAMP-CRP dependent, so their induction suggests a general shortage of preferred carbon sources rather than a response to maltose or glycerol specifically. Infection raises carbon demand across the culture, consistent with the bulk induction of de novo pyrimidine and UMP biosynthesis at 45 minutes (Table S2), and only cells with intact host transcription can mount such a catabolite-repression response, which would explain why the signature is confined to cells with low viral load.

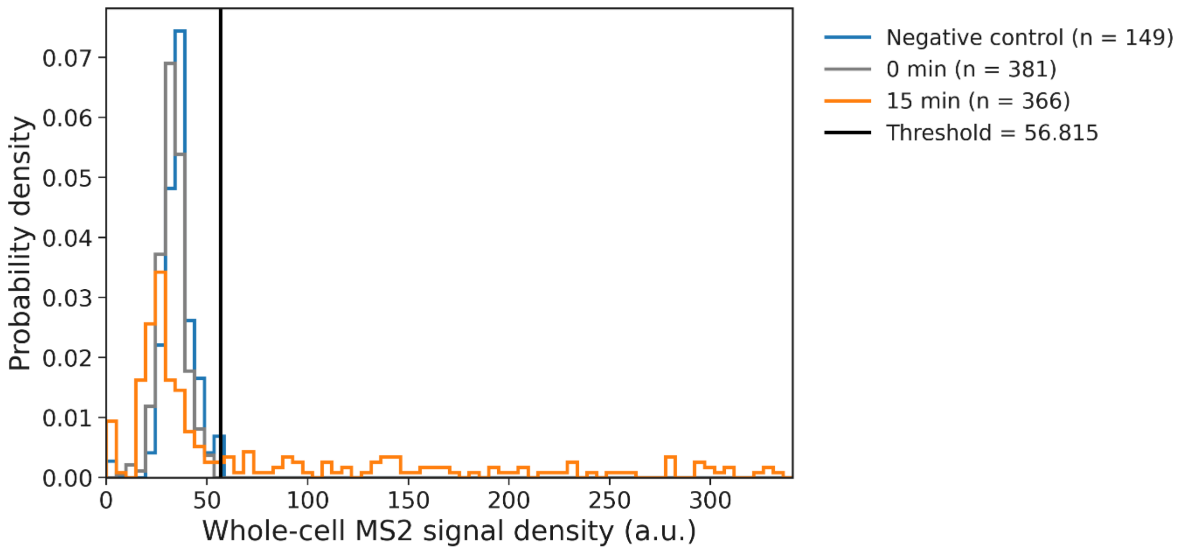


**Figure S10. smFISH MS2-positive threshold**. Distribution of whole-cell MS2 signal density for the negative control (no phage), an uninfected sample collected at the time of infection, and an infected 15-minute sample. Signal density is the background-corrected MS2 smFISH fluorescence integrated over the cell and divided by the cell area, in arbitrary units (a.u.) of camera intensity. The threshold for scoring a cell MS2-positive (black line) was set strictly above the maximum signal of the negative control. The 0-minute sample overlaps the negative control and lies entirely below the threshold, while the 15-minute sample shows a distinct positive population above it.

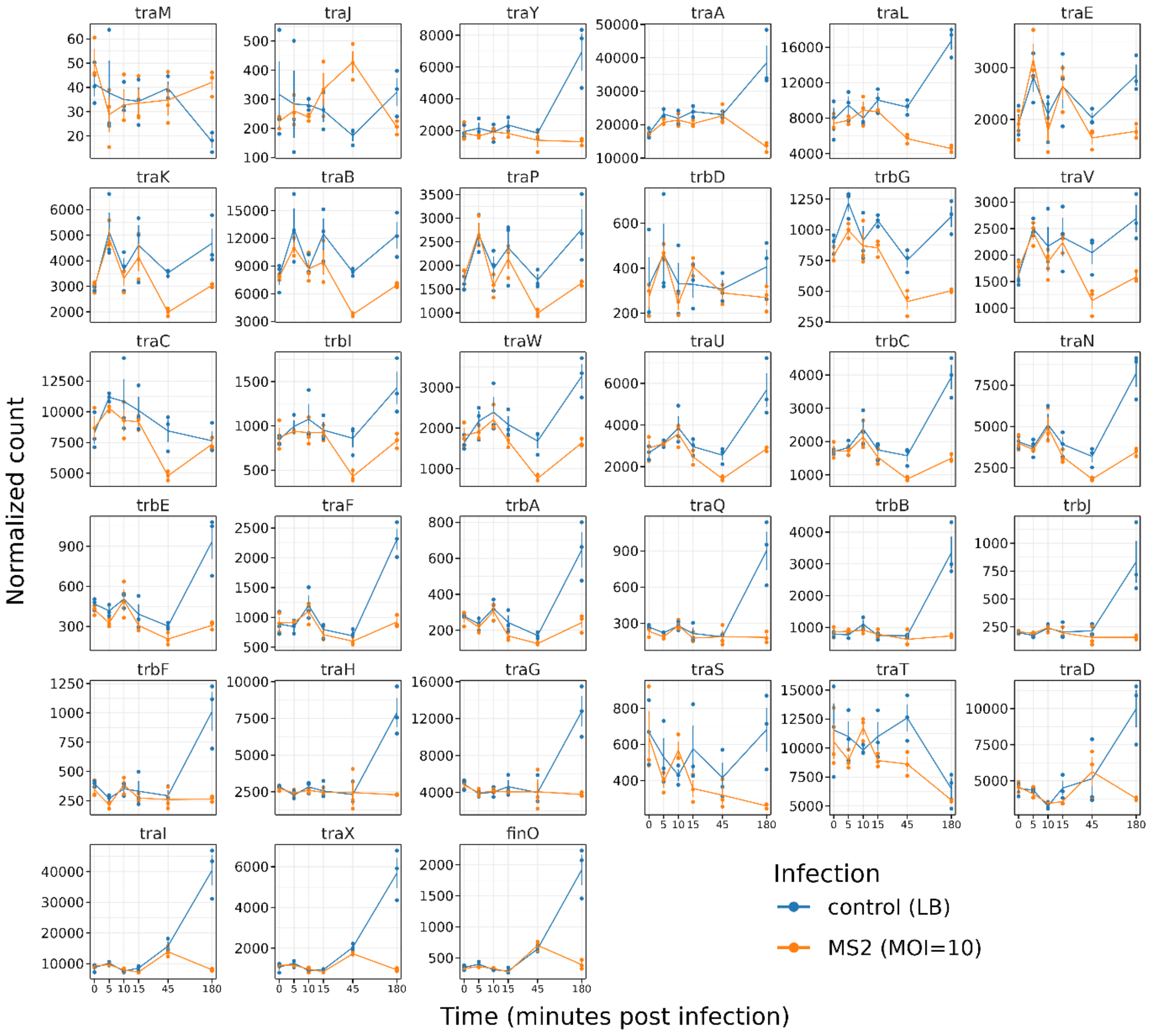


**Figure S11. Pilus-associated gene expression dynamics.** Normalized counts over the infection time course for 32 genes of the *tra* operon and the fertility inhibition regulator *finO*, in infected (orange) and time-matched uninfected control (blue) samples. Panels are ordered by gene position in the operon (left to right, top to bottom). Points show biological replicates and error bars the standard error. Most genes show lower expression in infected than in control samples at 180 minutes. This difference reflects both slightly reduced expression in the infected samples but mainly a marked increase in the controls at this time point. Expression in the infected samples at 180 minutes nonetheless remains comparable to, or only modestly below, that at earlier infected time points, indicating small transcriptional downregulation of the operon rather than its complete silencing.

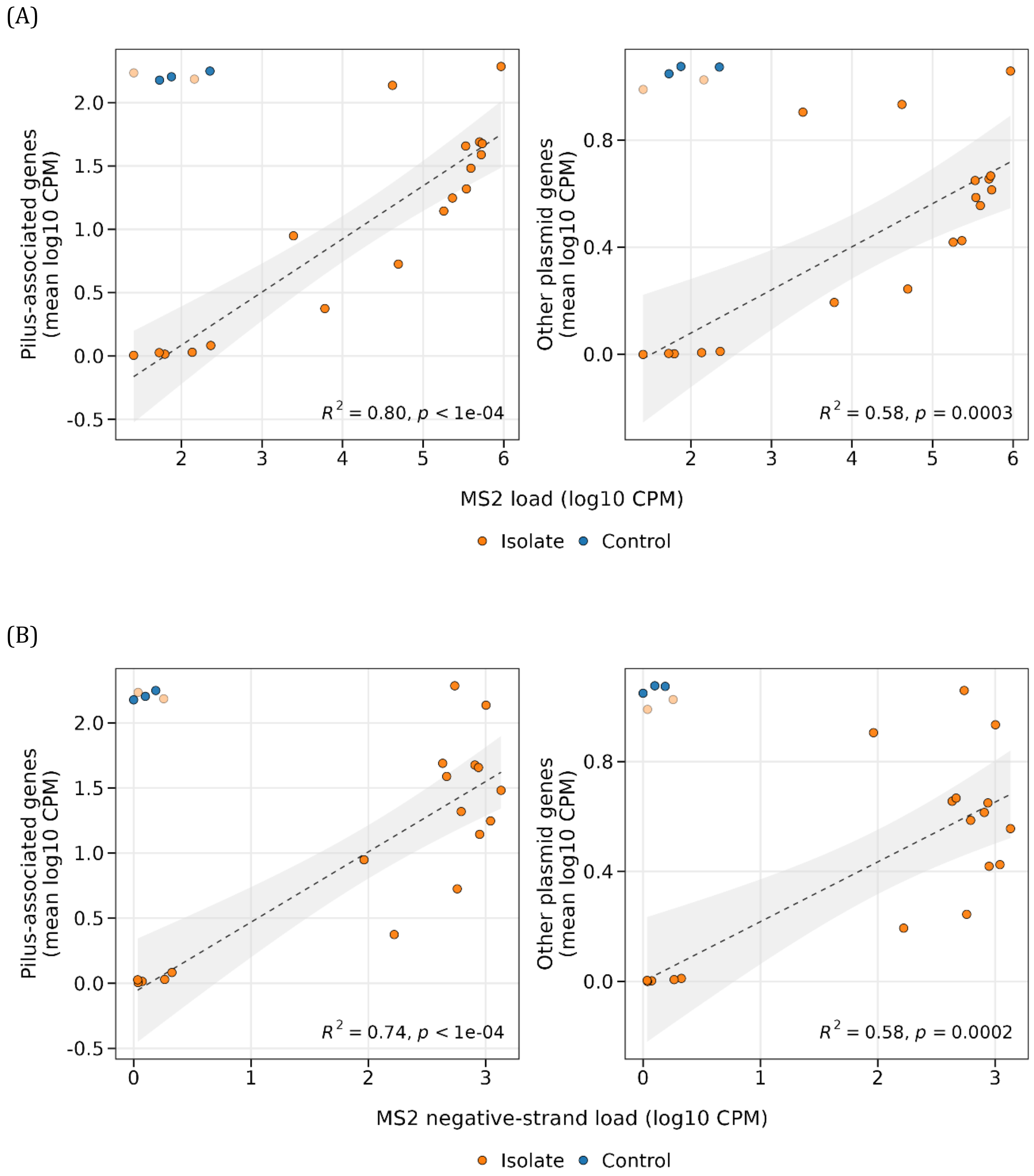


**Figure S12. Associations of pilus-associated and non-pilus F-plasmid genes with MS2 load in colony isolates.** (A) As in Fig. 4C, with F-plasmid genes partitioned into pilus-associated (n = 33, left) and non-pilus (n = 46, right) sets, plotted against positive-strand MS2 load (log10 CPM of host-mapped reads). (B) As in (A), with negative-strand MS2 RNA counts (log10 CPM of host-mapped reads) on the x-axis in place of positive-strand MS2. Negative-strand RNA is the MS2 replication intermediate and therefore reports productive infection. Lines show linear regressions across the 18 plaque-negative isolates with 95% confidence bands; control samples (blue) and two colony-derived revertants (light orange) were excluded from the regressions, as in Fig. 4C.

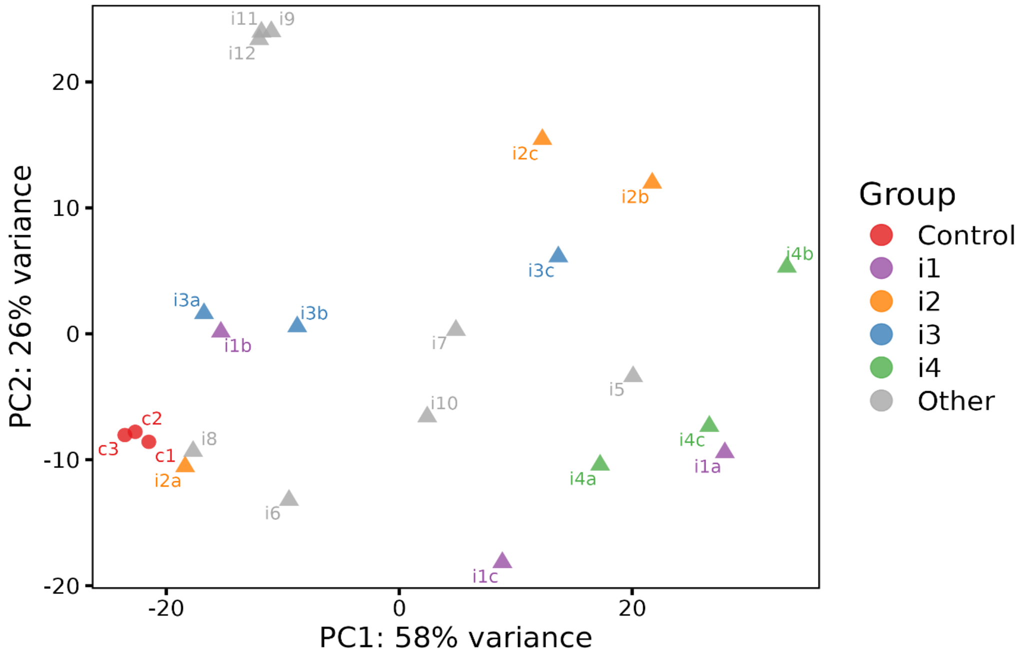


**Figure S13. PCA of transcriptomic profiles of phage-surviving isolates.** PCA was performed on the 500 most variable genes across all samples, using regularized log-transformed (rlog) counts. Each point is a sequencing library; the three uninfected control replicates are shown in red, the four isolates profiled in triplicate (i1–i4) in distinct colours, with replicates labelled a–c, and the remaining singly profiled isolates in grey. The control replicates cluster tightly, whereas the phage-exposed isolates are dispersed across both principal components, which together account for 84% of the variance. Notably, replicates of the same isolate are frequently as far apart as different isolates (for example i2a versus i2b and i2c, or i1b versus i1c), indicating that transcriptional variation arises between independent cultures of a single isolate and not only between isolates, consistent with the divergence expected between separate regrowths from a common stock (see also Table S6).

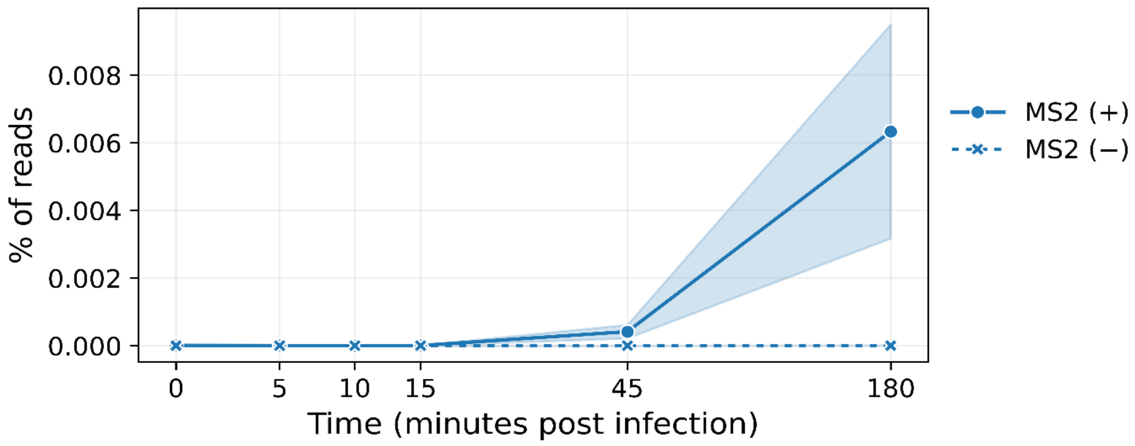


**Figure S14. Percent of MS2 reads over time in the control samples in bulk RNA seq.** Positive-strand (solid) and negative-strand (dashed) MS2 reads are shown as a percentage of mapped reads; lines are means of three biological replicates and shaded bands the standard error. Positive-strand MS2 remained below 0.01% of reads at all time points, while negative-strand MS2 was effectively absent (a single read across all 18 control samples). For comparison, MS2 accounted for approximately 74% and 77% of mapped reads in infected samples at 45 and 180 minutes post-infection.

**Supplementary tables**

**Table S1.** Genes significantly up- or downregulated (DESeq2, |log2 fold change| > 1.5, adjusted p < 0.05) in infected vs. control cells at each time point (0–180 min) and between time points within each condition. Available at Zenodo (https://doi.org/10.5281/zenodo.23232244)

**Table S2.** GO biological processes functional enrichment of gene list upregulated in infected population 45 minutes post infection (fold change > 1.5, adjusted p-value < 0.05). Similar GO terms were merged using String-DB merge rows when similarity >= 0.8.

| #term ID | term description | observed gene count | background gene count | strength | signal | false discovery rate |
|---|---|---|---|---|---|---|
| **GO:0006355** | Regulation of transcription, DNA-templated | 45 | 376 | 0.38 | 0.57 | 0.00027 |
| **GO:0050794** | Regulation of cellular process | 58 | 617 | 0.27 | 0.48 | 0.00059 |
| **GO:0006635** | Fatty acid beta-oxidation | 9 | 20 | 0.95 | 0.7 | 0.0013 |
| **GO:0045892** | Negative regulation of transcription, DNA-templated | 22 | 137 | 0.51 | 0.56 | 0.0013 |
| **GO:0046395** | Carboxylic acid catabolic process | 26 | 199 | 0.42 | 0.5 | 0.0020 |

| | | | | | | |
|---|---|---|---|---|---|---|
| **GO:0009220** | Pyrimidine ribonucleotide biosynthetic process | 7 | 13 | 1.03 | 0.65 | 0.0025 |
| **GO:0006631** | Fatty acid metabolic process | 14 | 70 | 0.6 | 0.54 | 0.0030 |
| **GO:0010124** | Phenylacetate catabolic process | 7 | 15 | 0.97 | 0.6 | 0.0038 |
| **GO:0006206** | Pyrimidine nucleobase metabolic process | 9 | 30 | 0.78 | 0.55 | 0.0047 |
| **GO:0048519** | Negative regulation of biological process | 28 | 245 | 0.36 | 0.43 | 0.0049 |
| **GO:0044205** | De novo UMP biosynthetic process | 5 | 7 | 1.15 | 0.54 | 0.0086 |
| **GO:0070689** | L-threonine catabolic process to propionate | 5 | 7 | 1.15 | 0.54 | 0.0086 |
| **GO:0072329** | Monocarboxylic acid catabolic process | 15 | 98 | 0.49 | 0.4 | 0.0144 |

| | | | | | | |
|---|---|---|---|---|---|---|
| **GO:0006054** | N-acetylneuraminate metabolic process | 5 | 9 | 1.05 | 0.45 | 0.0181 |
| **GO:0006207** | De novo pyrimidine nucleobase biosynthetic process | 5 | 9 | 1.05 | 0.45 | 0.0181 |

**Table S3. Number of cells profiled by smFISH across time points.**

| Experiment | Time point (min) | Fields | Cells | MS2-positive cells | MS2-positive (%) |
|---|---|---|---|---|---|
| Set 1 | Negative control | 10 | 149 | 0 | 0 |
| | 0 | 10 | 381 | 0 | 0 |
| | 15 | 10 | 366 | 208 | 56.8 |
| | 45 | 10 | 985 | 745 | 75.6 |
| | 120 | 24 | 217 | 182 | 83.9 |
| | 180 | 16 | 130 | 117 | 90 |
| | 240 | 11 | 234 | 190 | 81.2 |
| Set 2 | Negative control | 8 | 126 | 0 | 0 |
| | 0 | 7 | 165 | 1 | 0.6 |
| | 15 | 14 | 986 | 567 | 57.5 |
| | 45 | 13 | 805 | 655 | 81.4 |
| | 120 | 11 | 60 | 52 | 86.7 |
| | 180 | 7 | 36 | 30 | 83.3 |
| | 240 | 18 | 249 | 199 | 79.9 |

Of note, infection prevalence estimates from smFISH and scRNAseq are not directly comparable. smFISH scores every segmented object against a threshold derived from the uninfected control, and gave 76–81% positive at 45 min across technical replicates. scRNAseq scores only barcodes passing a host-RNA quality filter, and gave ~90–96% across filter settings (Fig. S7). Cells with low RNA content or incomplete permeabilisation are retained and scored negative in the former and excluded in the latter, which may account for the residual difference between the two methods. Set 2 is a technical replicate (the same samples re-imaged); Set 1 is shown in Fig. 2D and Fig. S10.

**Table S4.** GO biological processes functional enrichment of gene list upregulated in 180 minutes, specifically: gene list is composed of upregulated genes between 180 minutes and 15-minutes ms2-infected samples, removing genes upregulated180 minutes vs. 15 minutes in the control samples (fold change > 1.5, adjusted p value < 0.05). Similar GO terms were merged using String-DB merge rows when similarity >= 0.8. Notably, pathways related to biofilm formation are upregulated.

| #term ID | term description | observed gene count | background gene count | strength | signal | false discovery rate |
|---|---|---|---|---|---|---|
| **GO:0055085** | Transmembrane transport | 103 | 671 | 0.26 | 0.61 | 8.82E-06 |
| **GO:0098655** | Cation transmembrane transport | 49 | 223 | 0.41 | 0.74 | 8.82E-06 |
| **GO:0071705** | Nitrogen compound transport | 55 | 275 | 0.37 | 0.7 | 1.03E-05 |
| **GO:0006811** | Ion transport | 68 | 384 | 0.32 | 0.66 | 1.13E-05 |
| **GO:0071702** | Organic substance transport | 77 | 485 | 0.27 | 0.58 | 5.13E-05 |
| **GO:0030030** | Cell projection organization | 19 | 51 | 0.64 | 0.74 | 0.00019 |
| **GO:0071973** | Bacterial-type flagellum-dependent cell motility | 21 | 61 | 0.61 | 0.72 | 0.00019 |
| **GO:0044781** | Bacterial-type flagellum organization | 14 | 32 | 0.71 | 0.65 | 0.0011 |
| **GO:0006935** | Chemotaxis | 15 | 38 | 0.67 | 0.62 | 0.0013 |

| | | | | | | |
|---|---|---|---|---|---|---|
| **GO:0098660** | Inorganic ion transmembrane transport | 34 | 170 | 0.37 | 0.49 | 0.0020 |
| **GO:0009238** | Enterobactin metabolic process | 8 | 9 | 1.02 | 0.65 | 0.0024 |
| **GO:0046377** | Colanic acid metabolic process | 11 | 22 | 0.77 | 0.59 | 0.0028 |
| **GO:0006865** | Amino acid transport | 23 | 99 | 0.44 | 0.46 | 0.0053 |
| **GO:0006820** | Anion transport | 34 | 184 | 0.34 | 0.42 | 0.0057 |
| **GO:0007155** | Cell adhesion | 17 | 60 | 0.53 | 0.47 | 0.0062 |
| **GO:0140352** | Export from cell | 17 | 65 | 0.49 | 0.43 | 0.0101 |
| **GO:0044718** | Siderophore transmembrane transport | 8 | 14 | 0.83 | 0.48 | 0.0104 |
| **GO:0030031** | Cell projection assembly | 13 | 42 | 0.56 | 0.41 | 0.0143 |
| **GO:0098656** | Anion transmembrane transport | 27 | 145 | 0.34 | 0.35 | 0.0184 |

| | | | | | | |
|---|---|---|---|---|---|---|
| **GO:0000041** | Transition metal ion transport | 17 | 72 | 0.45 | 0.35 | 0.0241 |
| **GO:1902600** | Proton transmembrane transport | 17 | 72 | 0.45 | 0.35 | 0.0241 |
| **GO:0003333** | Amino acid transmembrane transport | 18 | 81 | 0.42 | 0.33 | 0.0291 |
| **GO:0043708** | Cell adhesion involved in biofilm formation | 13 | 47 | 0.51 | 0.34 | 0.0307 |
| **GO:0030254** | Protein secretion by the type III secretion system | 6 | 9 | 0.9 | 0.38 | 0.0312 |
| **GO:0006826** | Iron ion transport | 13 | 50 | 0.49 | 0.31 | 0.0443 |
| **GO:0045228** | Slime layer polysaccharide biosynthetic process | 6 | 10 | 0.85 | 0.34 | 0.0443 |

**Table S5. Whole-genome sequencing variants identified using LoFreq and Sniffles2.** The table lists all detected variants across all samples. Variants detected in all control samples were excluded as background relative to the reference genome. Of the remaining variants, those present exclusively in infected samples at a frequency > 0.95 were defined as mutations. Available at Zenodo (https://doi.org/10.5281/zenodo.23232244).

**Table S6. Mapping between DNA-seq and RNA-seq sample names.** Each isolate was preserved as a frozen stock, from which separate cultures were subsequently grown for DNA-seq and for RNA-seq; the two datasets therefore derive from independent regrowth of the same stock rather than from the same culture. Each isolate was sequenced at the DNA level once and is given a single DNA-seq identifier. Isolates for which RNA-seq was also performed carry a corresponding RNA-seq identifier; where an isolate was sequenced in triplicate, the replicates are distinguished by the suffixes a–c. Empty cells indicate isolates not included in the RNA-seq experiment. Variation in plasmid content, viral load, and sub-clonal genotypes may persist within a stock or arise and be selected during outgrowth, so genotypes detected in the DNA-seq culture are not necessarily present at the same frequency, or at all, in the corresponding RNA-seq culture described in the main text.

<table>
<tr><th>DNAseq sample name</th><th>RNAseq sample name</th><th>DNAseq mutations with frequency > 0.8<br>(gene, frequency, mutation)</th><th>Description</th></tr>
<tr><td>c1</td><td>c1</td><td></td><td rowspan="10">Uninfected control</td></tr>
<tr><td>c2</td><td>c2</td><td></td></tr>
<tr><td>c3</td><td>c3</td><td></td></tr>
<tr><td>c4</td><td></td><td></td></tr>
<tr><td>c5</td><td></td><td></td></tr>
<tr><td>c6</td><td></td><td></td></tr>
<tr><td>c7</td><td></td><td></td></tr>
<tr><td>c8</td><td></td><td></td></tr>
<tr><td>c9</td><td></td><td></td></tr>
<tr><td>c10</td><td></td><td></td></tr>
<tr><td rowspan="3">i1</td><td>i1a</td><td rowspan="3"></td><td rowspan="32">Exposed isolate</td></tr>
<tr><td>i1b</td></tr>
<tr><td>i1c</td></tr>
<tr><td rowspan="3">i2</td><td>i2a</td><td rowspan="3"></td></tr>
<tr><td>i2b</td></tr>
<tr><td>i2c</td></tr>
<tr><td rowspan="3">i3</td><td>i3a</td><td rowspan="3"></td></tr>
<tr><td>i3b</td></tr>
<tr><td>i3c</td></tr>
<tr><td rowspan="3">i4</td><td>i4a</td><td rowspan="3">wcaC, 0.96, C156G</td></tr>
<tr><td>i4b</td></tr>
<tr><td>i4c</td></tr>
<tr><td>i5</td><td></td><td></td></tr>
<tr><td>i6</td><td></td><td></td></tr>
<tr><td>i7</td><td></td><td></td></tr>
<tr><td>i8</td><td></td><td></td></tr>
<tr><td>i9</td><td></td><td></td></tr>
<tr><td>i10</td><td></td><td></td></tr>
<tr><td>i11</td><td></td><td></td></tr>
<tr><td>i12</td><td></td><td>ybaK, 0.99, R133R</td></tr>
<tr><td>i13</td><td></td><td></td></tr>
<tr><td>i14</td><td></td><td></td></tr>
<tr><td>i15</td><td></td><td>wcaD, 0.99, T201A</td></tr>
<tr><td>i16</td><td></td><td>yejK, 0.98, N295S</td></tr>
<tr><td>i17</td><td></td><td></td></tr>
<tr><td>i18</td><td></td><td></td></tr>
<tr><td>i19</td><td></td><td></td></tr>
<tr><td>i20</td><td></td><td></td></tr>
</table>

**Table S7. Probe set for the first scRNA-seq experiment.** Available at Zenodo (https://doi.org/10.5281/zenodo.23232244).

**Table S8. Second scRNAseq probe table.** All probes from Table S7 were included, along with the additional probes listed here, which served as feature barcodes. The MS2 probes from Table S7 were omitted, as they were instead detected as feature barcodes. Available at Zenodo (https://doi.org/10.5281/zenodo.23232244).
